# Heritability enrichment in context-specific regulatory networks improves phenotype-relevant tissue identification

**DOI:** 10.1101/2022.09.06.506769

**Authors:** Zhanying Feng, Zhana Duren, Jingxue Xin, Qiuyue Yuan, Yaoxi He, Bing Su, Wing Hung Wong, Yong Wang

## Abstract

Systems genetics holds the promise to decipher complex traits by interpreting their associated SNPs through gene regulatory networks derived from comprehensive multi-omics data of cell types, tissues, and organs. Here, we propose SpecVar to integrate paired chromatin accessibility and gene expression data into context-specific regulatory network atlas and regulatory categories, conduct heritability enrichment analysis with GWAS summary statistics, identify relevant tissues, and depict shared heritability and regulations by relevance correlation. Our method improves power upon existing approaches by associating SNPs with context-specific regulatory elements to assess heritability enrichments and by explicitly prioritizing gene regulations underlying relevant tissues. Experiments on GWAS of six phenotypes show that SpecVar can improve heritability enrichment, accurately detect relevant tissues, and reveal causal regulations. Furthermore, SpecVar correlates the relevance patterns for pairs of phenotypes and better reveals shared heritability and regulations of phenotypes than existing methods. Studying GWAS of 206 phenotypes in UK-Biobank demonstrates that SpecVar leverages the context-specific regulatory network atlas to prioritize phenotypes’ relevant tissues and shared heritability for biological and therapeutic insights. SpecVar provides a powerful way to interpret SNPs via context-specific regulatory networks and is available at https://github.com/AMSSwanglab/SpecVar.

## Introduction

Genome-Wide Association Studies (GWAS) have gained a great success to identify thousands of genetic variants significantly associated with a variety of human complex phenotypes. Interpretation of those genetic variants holds the key to biological mechanism discovery and personalized medicine practice. However, this task is hindered by the genetic architecture that the heritability is distributed across SNPs of the whole genome with linkage disequilibrium (LD), cumulatively affecting complex traits. By quantifying the contribution of true polygenic signal considering linkage disequilibrium, LD Score regression (LDSC) provides a widely appreciated method to estimate heritability (B. K. Bulik-Sullivan et al., 2015) and genetic correlation (B. Bulik-Sullivan et al., 2015) from GWAS summary statistics.

Another obstacle to genetic variant interpretation is that SNPs contribute to phenotype through gene regulatory networks in certain cellular contexts, i.e., causal tissues or cell types. Those tissues are characterized by different types of epigenetic data, which give the active regions of the genome that interact with transcription factors (TF) to regulate gene expression. Stratified LDSC (S-LDSC) extends LDSC and can estimate the partitioned heritability enrichment in the functional categories (Finucane et al., 2015). The categories can be non-specific genome annotations (such as coding, UTR, promoter, and intronic regions) and context-specific regulatory regions called from chromatin data of different cell types, such as DNase-I hypersensitive sites from DNase-seq data, accessible peaks from ATAC-seq data, histone marker or TF binding sites from ChIP-seq data (LDSC-AAP and LDSC-SAP). Using expression data, the functional categories can be alternatively constructed by the 100-kb windows around the transcribed regions of specifically expressed genes (LDSC-SEG) (Hilary K. Finucane et al., 2018). Essentially, these strategies summarize the high dimensional SNP signals from the whole genome into partitioned heritability enrichments and successfully identify relevant cellular tissues for many phenotypes (Finucane et al., 2015).

The rapid increase of multi-modal data resources, especially matched gene expression, chromatin states, and TF binding sites (i.e., measured on the same sample), offers an exciting opportunity to construct better functional categories for estimating heritability enrichment. One efficient way is to integrate large-scale epigenomic and transcriptomic data spanning diverse human contexts to infer regulatory networks (Duren et al., 2017). Those regulatory networks provide rich context-specific information and usually comprise TFs, regulatory elements (REs), and target genes (TGs). Recently, we developed the PECA2 model to infer regulatory network from paired expression and chromatin accessibility data (Duren et al., 2017; Duren et al., 2020). The inferred regulatory networks have been used to identify the master regulators in stem cell differentiation (Li et al., 2019) and to interpret conserved regions for the non-model organisms (Xin et al., 2020). Non-coding genetic variants can be interpreted in the regulatory networks on how they cooperatively affect complex traits through gene regulation in certain tissues or cell types. For example, genetic variants in the regulatory network of cranial neural crest cells are elucidated on how they affect human facial morphology (Feng et al., 2021). RSS-NET utilizes gene regulatory networks of multiple contexts and shows better tissue enrichment estimation by decomposing the total effect of a SNP through TF-TG regulations (Zhu et al., 2021) and HiChIP RE-TG regulations (Ma et al., 2022). And the phenotype-associated SNPs often function in a tissue- or cell-type-specific manner (Westra & Franke, 2014). The advances in constructing regulatory networks and interpreting genetic variants with regulatory networks enlighten us to 1) assemble a more comprehensive context-specific regulatory network atlas by using paired expression and accessibility data across diverse cellular contexts; 2) build context-specific regulatory categories by focusing on RE’s specificity of regulatory networks; 3) systematically identify enriched tissues or cell types, shared heritability (van Rheenen et al., 2019), and the underlying gene regulations of phenotypes.

Specifically, we proposed SpecVar to first leverage the publicly available paired expression and chromatin accessibility data in ENCODE and ROADMAP to systematically construct context-specific regulatory networks of 77 human contexts, covering major cell types and germ layer lineages. This atlas served as a valuable resource for genetic variants interpretation in multi-cellular contexts. SpecVar then used this atlas to construct regulatory categories in the genome. The heritability enrichment of GWAS was shown to be significantly improved by our context-specific regulatory categories. Based on the heritability enrichment and P-value in our regulatory categories, SpecVar defined the relevance score to give the context-specific representation of the GWAS. We showed that, for a single phenotype, the relevance score of SpecVar could identify relevant tissues more efficiently; and for multiple phenotypes, SpecVar could use relevance correlation to reveal shared heritability, common relevant tissues, and underlying gene regulations. These results showed that SpecVar is promising to serve as a tool for post-GWAS analysis.

## Results

### Overview of SpecVar method

SpecVar assembled a context-specific regulatory network atlas and built the context-specific representation (relevance score and SNP-associated regulatory network) of GWAS summary statistics based on heritability enrichment. **Fig. 1** summarized the major steps of SpecVar to construct context-specific regulatory network atlas and regulatory categories, calculate heritability enrichment and SNP-associated regulatory network, and investigate interpretable relevant tissues and relevance correlation.

**Fig. 1.**
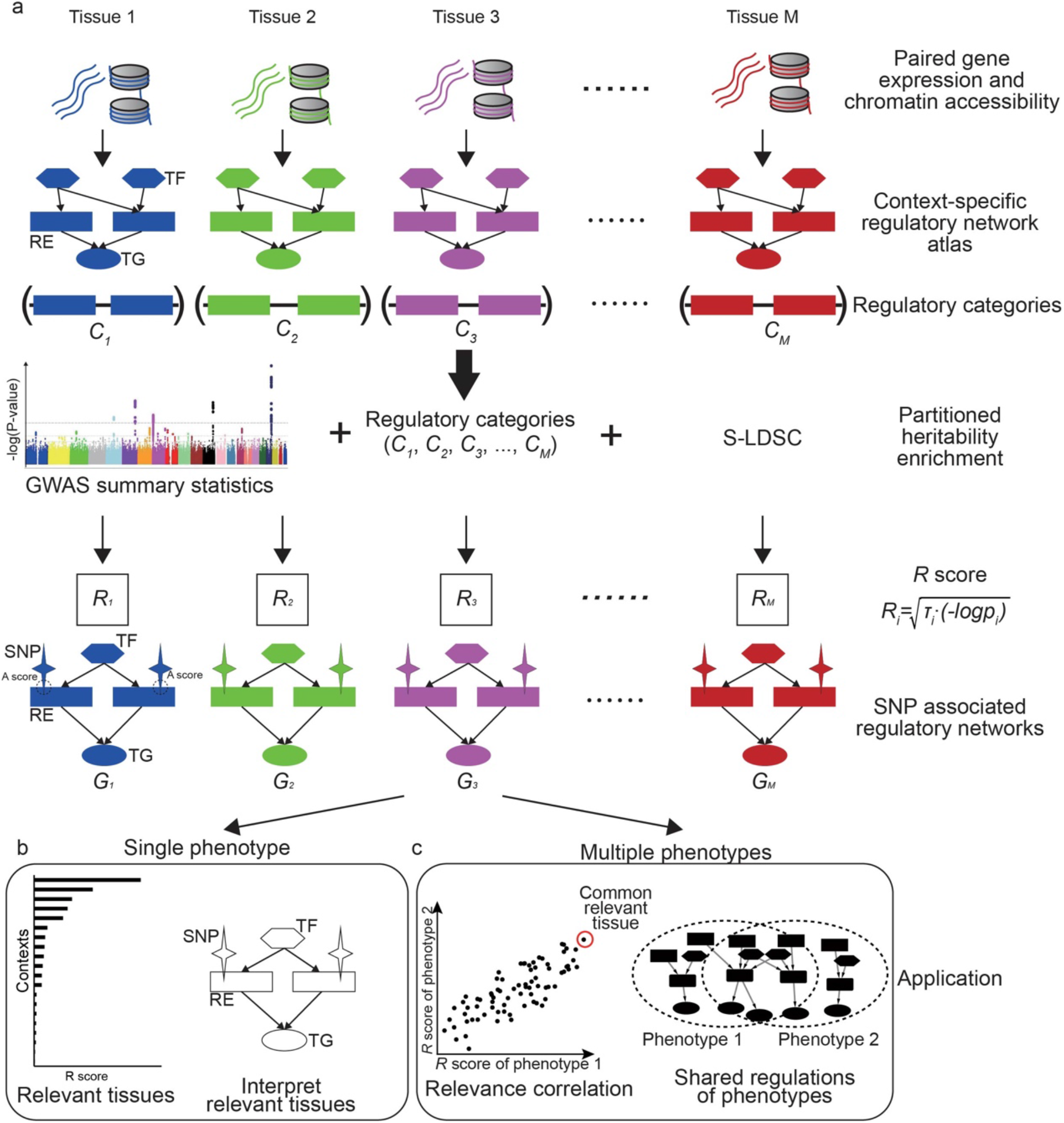
Overview of SpecVar. (a) SpecVar constructs an atlas of context-specific regulatory networks and regulatory categories. Then SpecVar represents GWAS summary statistics into relevance score and SNP-associated regulatory subnetworks. (b) For a single phenotype, SpecVar can use relevance score and SNP-associated regulatory subnetworks to identify and interpret relevant tissues. (c) For multiple phenotypes, based on relevance score, SpecVar can reveal relevance correlation, common relevant tissues, and shared regulations.

We first reconstructed regulatory networks of *M* (*M*=77 in this paper) contexts. Each network is represented by a set of relations between TF and RE and between RE and TG. The *M* contexts included samples from all three germ layers, such as “frontal cortex” (ectoderm), “fetal thymus” (mesoderm), and “body of pancreas” (endoderm), which ensured the wide coverage and system-level enrichment (**Fig. S1**). The context-specific regulatory networks were extracted based on the specificity of REs in each context’s regulatory network compared to other contexts, considering the hierarchical relationship of *M* contexts (**Methods, Table S1**). The REs in the *i*-th context-specific regulatory network were pooled to form a regulatory category *C*_*i*_ in the genome, which restricted the annotation to context-specific REs associated with active binding TFs and nearby regulated TGs (**Fig. 1a**). Our atlas leads to *M* regulatory categories, *C*_1_, *C*_2_, …, *C*_*M*_ of SpecVar. Given GWAS summary statistics, the *M* regulatory categories allowed partitioned heritability enrichment analysis by S-LDSC. For a phenotype, S-LDSC modeled genome-wide polygenic signal, partitioned SNPs into categories with different contributions for heritability, and considered SNP’s linkage disequilibrium with the following polygenic model:

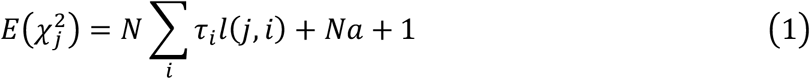

Here 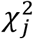 was the marginal association of SNP *j* from GWAS summary statistics; *N* was the sample size; 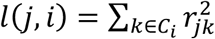 was the LD score of SNP *j* in the *i*-th regulatory category *C*_*i*_, where *r*_*jk*_ was the correlation between SNP *j* and SNP *k* in population; *a* measured the contribution of confounding biases; and *τ*_*i*_ represented the heritability enrichment of SNPs in *C*_*i*_. S-LDSC estimated the P-value *p*_*i*_ for the heritability enrichment (Finucane et al., 2015).

We defined the relevance score (*R*_*i*_) of this phenotype to *i*-th context (**Fig. 1a**) as follows by combining the enrichment score and statistical significance (P-value):

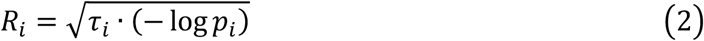

The relevance score (*R* score) provided a decision trade-off between the heritability enrichment and P-value resulting from a hypothesis test. It offered a new robust means to rank and select relevant tissues for a given phenotype (Xiao et al., 2014).

Meanwhile, SpecVar associated SNPs with context-specific regulatory networks for biological interpretation. We defined an association score (*A* score) to prioritize the REs by combining its regulatory strength and association significance with the phenotype (averaged −*logP* of SNPs located near the RE and down-weighted by their LD scores and distance to this RE). We extracted the REs with significant *A* scores (*P* ≤ 0.05), as well as their directly linked upstream TFs, downstream TGs, and associated SNPs, to form the SNP-associated regulatory subnetwork (**Fig. 1a, Methods**). Given GWAS summary statistics of a phenotype, SpecVar obtained *M* SNP-associated regulatory subnetworks, *G*_1_, *G*_2_, …, *G*_*M*_, allowing to interpret relevant tissues by SNP’s regulation mechanism.

The relevance score to diverse human contexts and SNP-associated regulatory networks allowed SpecVar to perform post-GWAS analysis. For a single phenotype, the *R* scores indicated the relevance of this phenotype to *M* contexts, which could be used to identify relevant tissues. Then in the relevant tissues, we could investigate the SNP-associated regulatory subnetwork to interpret the relevance (**Fig. 1b, Methods**). For multiple phenotypes, we could correlate the *R* score vectors in multiple contexts to define relevance correlation (Hilary K. Finucane et al., 2018). The relevance correlation might give insights into the association of phenotypes since SpecVar could further interpret the relevance correlation between two phenotypes by common relevant tissues and the overlapped SNP-associated regulatory subnetwork in common relevant tissues (**Fig. 1c, Methods**).

### Context-specific regulatory networks improve heritability enrichment

We first designed experiments to show that the context-specific regulatory networks could improve heritability enrichment. We collected GWAS summary statistics of six phenotypes, including two lipid phenotypes (Willer et al., 2013): low-density lipoprotein (LDL) and total cholesterol (TC); two human intelligential phenotypes (Lee et al., 2018): educational attainment (EA) and cognitive performance (CP); and two craniofacial bone phenotypes: brain shape (Naqvi et al., 2021) (BrainShape) and facial landmark point distances (Xiong et al., 2019) (Face). We used these six phenotypes as a benchmark since their relevant tissues have been studied and partially known: lipid phenotypes are associated with the liver for its key role in lipid metabolism (Nguyen et al., 2008); human intelligential phenotypes are associated with brain tissues (Goriounova & Mansvelder, 2019); Face and BrainShape had shared heritability in cranial neural crest cells (CNCC) (Naqvi et al., 2021). We compared our context-specific regulatory networks with four alternative methods of functional categories: all regulatory elements (ARE), all accessible peaks (AAP), specifically accessible peaks (SAP) (Finucane et al., 2015), and specifically expressed genes (Hilary K. Finucane et al., 2018) (SEG) (**Methods**).

First, we showed that SpecVar could achieve higher heritability enrichment in the relevant tissues than other methods. For LDL and TC, SpecVar obtained the highest heritability in their relevant tissue “right lobe of liver” than the other four methods (**Fig. 2a, b**). For EA and CP, they were relevant to brain tissues: “frontal cortex”, “cerebellum”, “caudate nucleus”, “Ammon’s horn” and “putamen”. SpecVar obtained the highest averaged heritability enrichment in these five brain tissues than the other four methods (**Fig. 2c, d**). For BrainShape and Face, SpecVar obtained a higher heritability enrichment in their relevant context “CNCC” than the other four methods (**Fig. 2e, f**). Second, except for the known relevant tissues, these complex traits may be relevant to other contexts. So, for every method, we ranked the heritability enrichment to get the top 10 contexts and used the top contexts’ heritability enrichment to compare the ability of these five methods to explain heritability in certain tissues or cell types. SpecVar also showed the best performance of heritability enrichment among the fiver methods (**Fig. 2g**). Taking BrainShape for example, SpecVar achieved significantly higher heritability enrichment (averaged heritability enrichment 96.13) than LDSC-ARE (26.77, *P* = 3.42 × 10^−3^), LDSC-SAP (42.92, *P* = 1.85 × 10^−2^), LDSC-SAP (20.34, *P* = 1.84 × 10^−3^), and LDSC-SEG (2.25, *P* = 3.05 × 10^−4^). We found specificity could significantly improve the heritability enrichment. Among the five methods in our comparison, SpecVar and LDSC-SAP are categories based on the specificity of LDSC-ARE and LDSC-AAP, respectively. SpecVar showed significantly higher heritability enrichment than LDSC-ARE and LDSC-SAP showed significantly higher heritability enrichment than LDSC-AAP (**Fig. 2g**). For BrainShape, SpecVar obtained averaged heritability enrichment of 96.31 of the top 10 contexts, which was significantly higher than LDSC-ARE (averaged heritability enrichment 26.77, *P* = 3.42 × 10^−3^); LDSC-SAP obtained average heritability enrichment of 42.92, and LDSC-AAP’s averaged heritability enrichment was 20.34 (*P* = 2.68 × 10^−3^). The other five phenotypes showed a similar improvement (**Fig. 2g**).

**Fig. 2.**
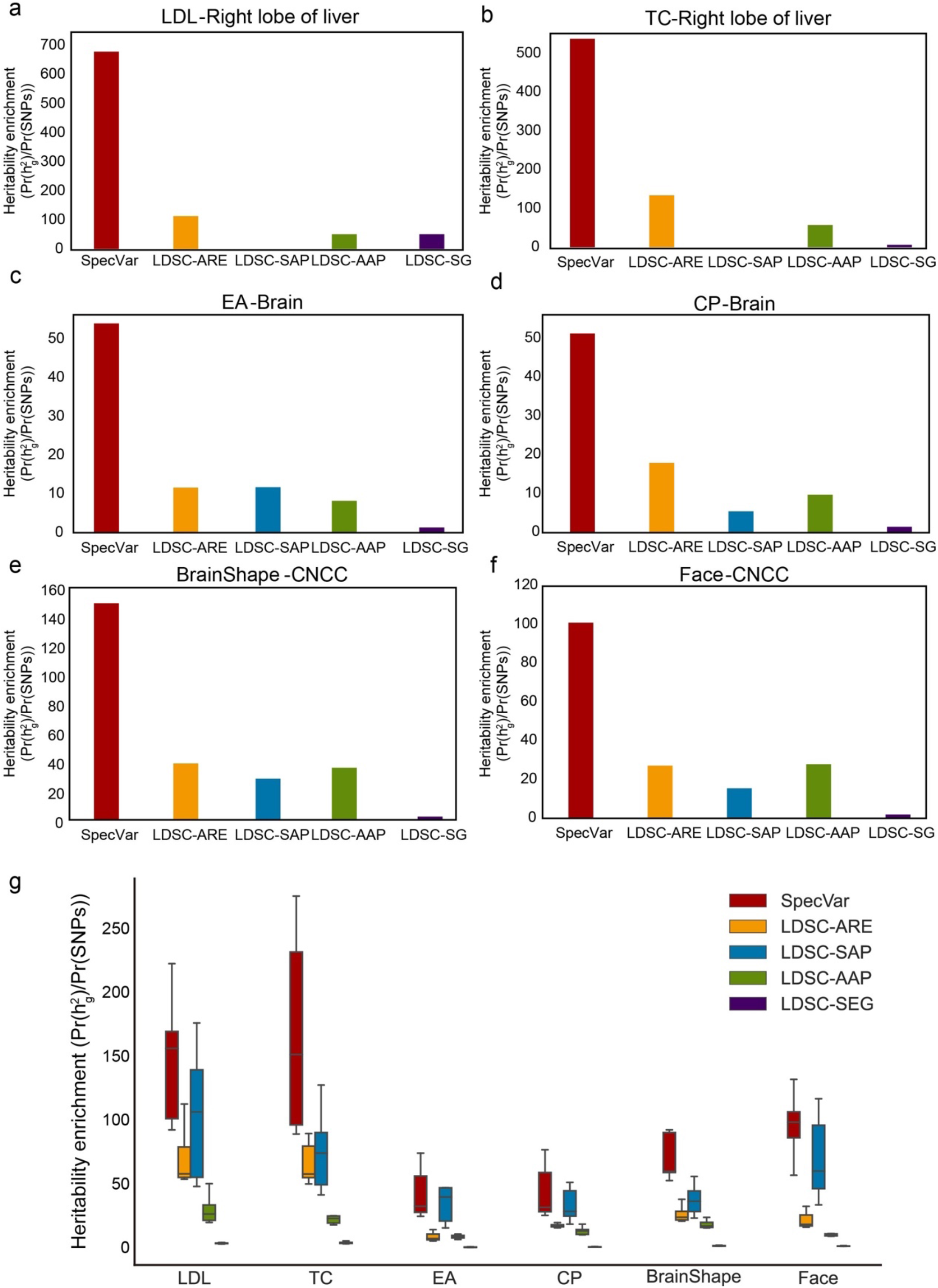
(a) The heritability enrichment of LDL in the “right lobe of liver” by five regulatory categories methods. (b) The heritability enrichment of TC in the “right lobe of liver” by five regulatory categories methods. (c) The five brain tissues’ averaged heritability enrichment of EA by five regulatory categories methods. (d) The five brain tissues’ averaged heritability enrichment of CP by five regulatory categories methods. (e) The heritability enrichment of BrainShape in “CNCC” by five regulatory categories methods. (f) The heritability enrichment of Face in “CNCC” by five regulatory categories methods. (g) Boxplot of top 10 tissues’ heritability enrichment for each of the five regulatory categories.

In summary, the experiment on six phenotypes’ GWAS summary statistics proved that SpecVar achieved the best performance in explaining the heritability of phenotypes. This demonstrated the power of integrating expression and chromatin accessibility data and considering contexts’ specificity.

### SpecVar can accurately reveal relevant tissues for phenotypes

After establishing that SpecVar could use the context-specific regulatory networks to improve heritability enrichment, we next showed that for given phenotype, SpecVar could use *R* scores identify relevant tissues more accurately than other methods of functional categories. In this experiment, we also used the above six phenotypes with their known relevant tissues as a benchmark and compared SpecVar to the other two specificity-based methods: LDSC-SAP and LDSC-SEG (**Methods**).

For two lipid phenotypes, SpecVar revealed that both LDL and TC were most significantly relevant to the “right lobe of liver” (**Fig. 3a, b, Table 1**), which was consistent with the existing reports that the liver plays a central role in lipid metabolism, serving as the center for lipoprotein uptake, formation, and export to the circulation (Jha et al., 2018; Nguyen et al., 2008). SpecVar found TC was significantly relevant to the “fetal adrenal gland” and the adrenal cortex has been revealed to play an important role in lipid mentalism (Boyd et al., 1983). However, LDSC-SAP and LDSC-SEG failed to prioritize liver tissue as the significant relevant tissue. For LDL, LDSC-SAP identified the “frontal cortex” to be the most relevant tissue. LDSC-SEG identified the most relevant tissue to be “HepG2”, which was human hepatoma cell lines, but the relevance score was not significant (**Fig. 3a, Table S2**). For TC, LDSC-SAP identified the “fetal adrenal gland” and LDSC-SEG obtained “HepG2” with an insignificant relevance score (**Figure 3b, Table S2**).

**Fig. 3.**
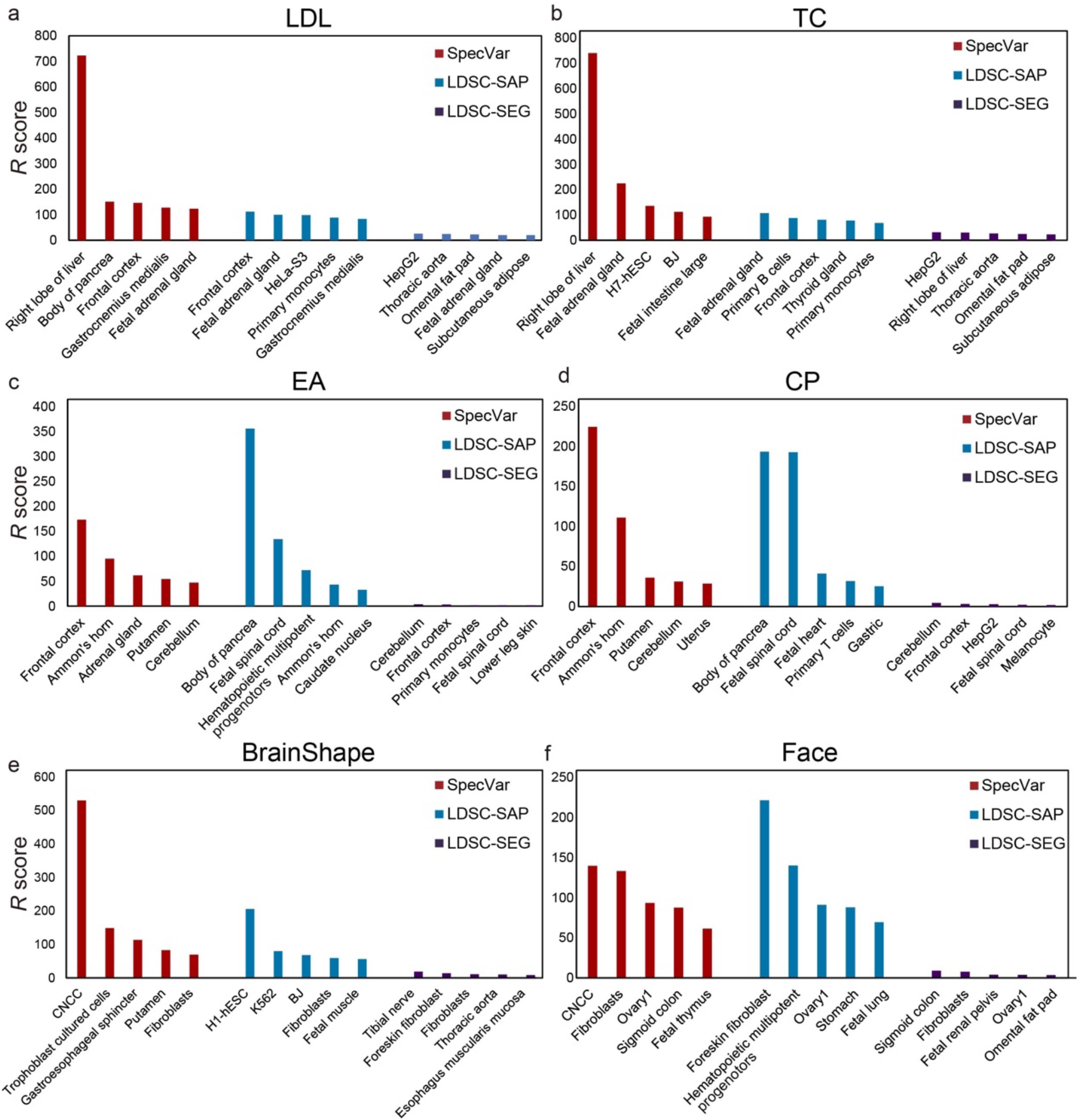
The top 5 relevant tissues ranked by the relevant score of SpecVar, LDSC-SAP, and LDSC-SEG for (a) LDL, (b) TC, (c) EA, (d) CP, (e) BrainShape, and (f) Face. Compared to LDSC-SAP and LDSC-SEG, SpecVar identified relevant tissue more accurately and stably.

**Table 1.**
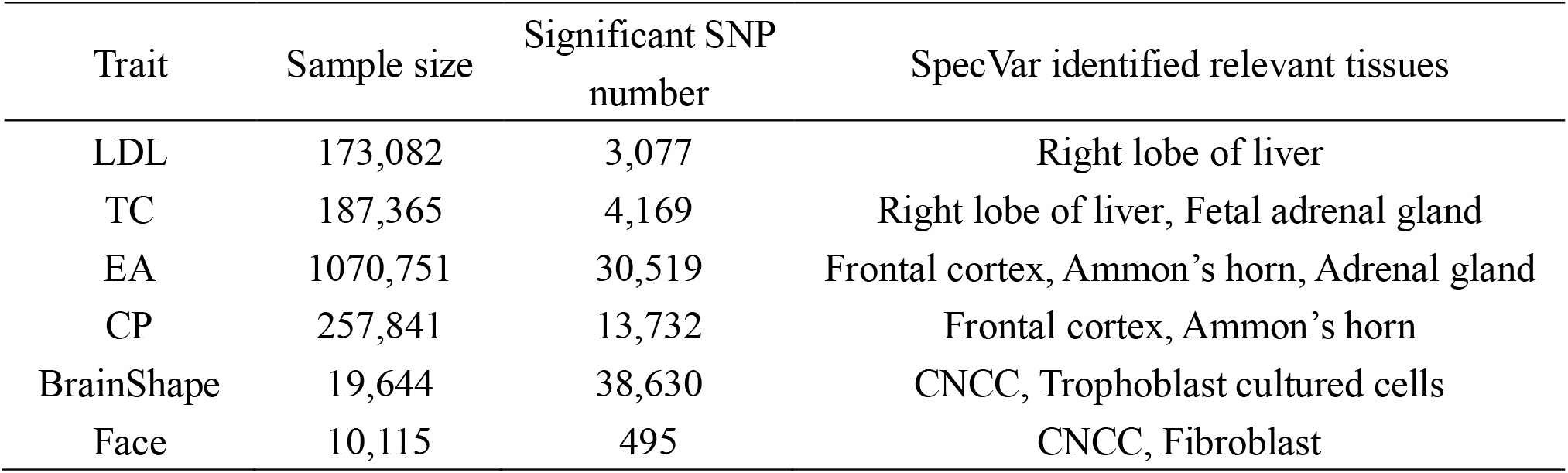
The total sample size, number of significant SNPs, and SpecVar identified relevant tissues of six phenotypes.

| Trait | Sample size | Significant SNP number | SpecVar identified relevant tissues |
| --- | --- | --- | --- |
| LDL | 173,082 | 3,077 | Right lobe of liver |
| TC | 187,365 | 4,169 | Right lobe of liver, Fetal adrenal gland |
| EA | 1070,751 | 30,519 | Frontal cortex, Ammon’s horn, Adrenal gland |
| CP | 257,841 | 13,732 | Frontal cortex, Ammon’s horn |
| BrainShape | 19,644 | 38,630 | CNCC, Trophoblast cultured cells |
| Face | 10,115 | 495 | CNCC, Fibroblast |

For two human intelligential phenotypes, SpecVar prioritized the “frontal cortex” to be the most relevant tissue for both EA and CP (**Fig. 3c, d, Table 1**). “Frontal cortex” is the cerebral cortex covering the front part of the frontal lobe and is implicated in planning complex cognitive behavior, personality expression, decision making, and moderating social behavior (Gabrieli et al., 1998; Yang & Raine, 2009). There were five tissues (“frontal cortex”, “Ammon’s horn”, “cerebellum”, “putamen”, “caudate nucleus”) from the brain in our atlas and they were significantly higher ranked by SpecVar’s relevance score than non-brain tissues for EA (Wilcoxon Rank-Sum test, *P* = 6.07 × 10^−7^, **Fig. 3c**) and CP (*P* = 8.00 × 10^−6^, **Fig. 3d**). In comparison, for EA, LDSC-SAP prioritized brain tissues to be higher ranked than non-brain tissues, but with a less significant P-value (*P* = 2.28 × 10^−3^, **Fig. 3c, Table S2**). LDSC-SEG could not rank brain tissues to be higher than non-brain tissues (*P* = 0.64, **Fig. 3c, Table S2**). For CP, LDSC-SAP failed to rank brain tissues as the most relevant tissues (*P* = 0.06, **Fig. 3d, Table S2**), and LDSC-SEG identified brain tissues to be more relevant than non-brain tissues but with a less significant P-value (*P* = 3.18 × 10^−3^, **Fig. 3d, Table S2**).

For both Face and BrainShape, SpecVar identified cranial neural crest cell (CNCC) as the most relevant context (**Fig. 3e, f, Table 1**). CNCC is a migratory cell population in early human craniofacial development that gives rise to the peripheral nervous system and many non-neural tissues such as smooth muscle cells, pigment cells of the skin, and craniofacial bones, which make it much more related to facial morphology and brain shape than the other 76 contexts (Cordero et al., 2011; “Neural crest makes a face,” 2008). Face morphology and brain shape were also revealed to share heritability in CNCC (Naqvi et al., 2021). But the other two methods failed to identify CNCC as the most relevant context. For BrainShape, LDSC-SAP identified “H1-hESC” and LDSC-SEG identified “tibial nerve” to be the most relevant tissue (**Fig. 3e, Table S2**). For Face, LDSC-SAP and LDSC-SEG identified “foreskin” and “sigmoid colon” to be the most relevant tissues, respectively (**Fig. 3f, Table S2**).

After identifying the relevant tissues, SpecVar could further interpret the relevance by extracting SNP-associated regulatory subnetwork (**Methods**). For example, we obtained BrainShape’s SNP-associated regulatory subnetwork in CNCC (**Fig. 4a**). There were 62 SNPs associated with 24 REs, 73 TFs, and 52 TGs. The TGs were tightly involved with brain development. For example, *POU3F3* is a well-known transcription factor involved in the development of the central nervous system and is related to many neurodevelopmental disorders (Blok et al., 2019). *EMX2* is expressed in the developing cerebral cortex and involved in the patterning of the rostral brain (Cecchi & Boncinelli, 2000). *FOXC2* is a member of the FOX family, which were modular competency factors for facial cartilage (Xu et al., 2018), and its mutation is linked to the cleft palate (Bahuau et al., 2002). By GWAS study, *FOXC2* was previously found to be associated with brain shape by its nearest significant SNP “16:86714715” (Naqvi et al., 2021). However, in CNCC, we did not find any accessible peaks that overlapped with this SNP. Instead, we found a CNCC-specific RE that regulated *FOXC2* in a locus of the 650k downstream. GWAS revealed the SNPs in this region had a strong association with brain shape and had high LD with each other (**Fig. 4b**). Our CNCC-specific regulations further prioritized only two SNPs (“16:87237097”, “16:87236947”) located in this CNCC-specific RE, which may influence the expression of *FOXC2* and the brain shape phenotypes. This example showed the power of SpecVar to interpret the genetic variants’ association to phenotypes with detailed regulatory networks in relevant tissues.

**Fig. 4.**
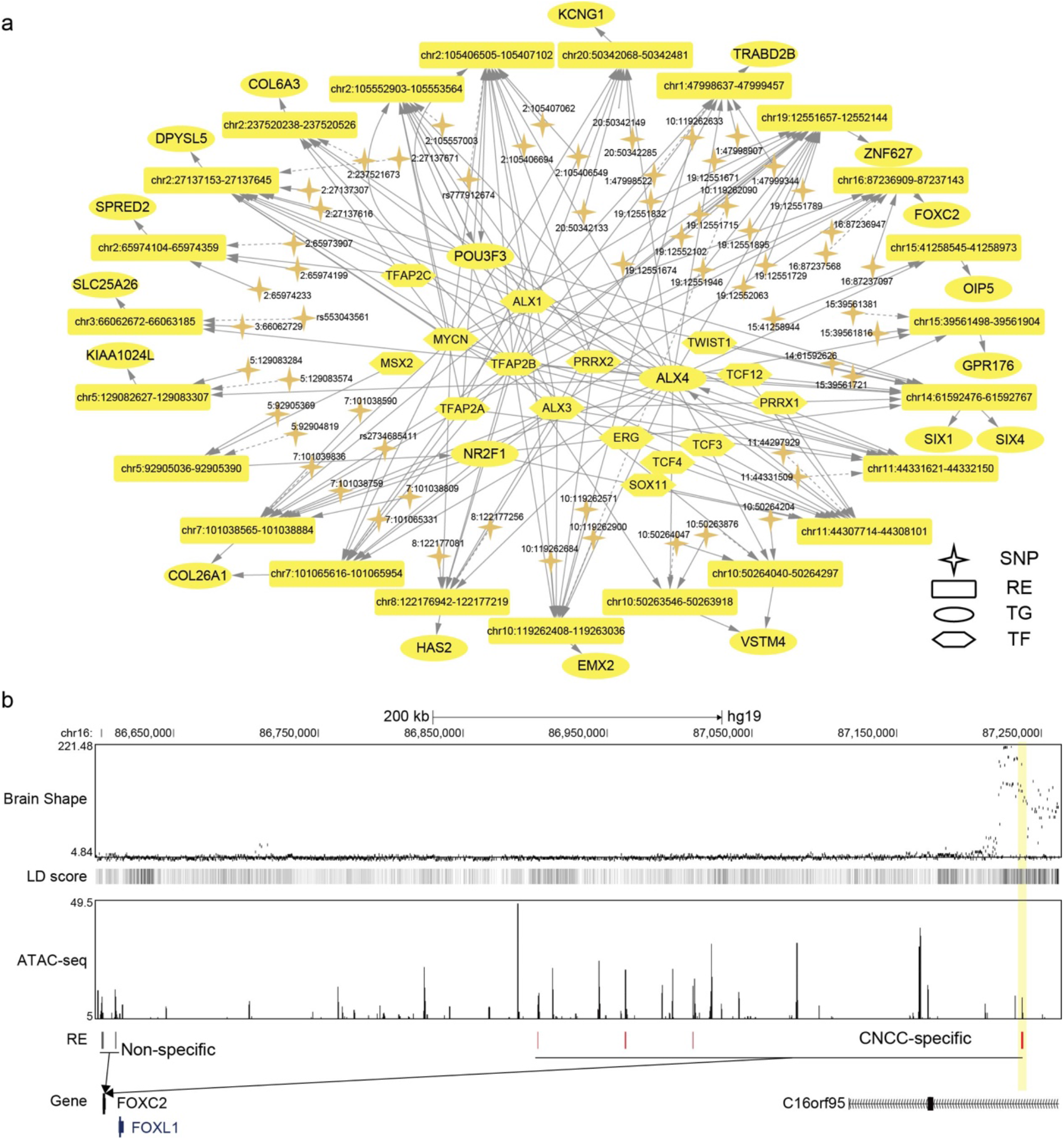
(a) The BrainShape’s SNP-associated regulatory subnetwork in CNCC. The dash arrows indicate significant SNPs that are not located in RE but near this RE. (b) SNP associated regulation of *FOXC2*. There is a group of significant SNPs of BrainShape that is located in the 650k downstream of *FOXC2* and they are with high linkage disequilibrium. SpecVar prioritizes SNPs located in a CNCC-specific RE as causal genetic variants affecting brain shape through regulation of *FOXC2*.

In summary, we evaluated SpecVar’s ability to identify relevant tissues using six well-studied phenotypes as the gold standard by comparison with the functional categories of LDSC-SAP and LDSC-SEG. The results showed that SpecVar could identify relevant tissues more accurately and stably and meanwhile provide detailed regulations to interpret the relevance to tissues.

### SpecVar reveals the association of multiple phenotypes by relevance correlation

SpecVar’s accurate and robust relevance to tissues enlightens us to define the relevance correlation of two phenotypes by Spearman correlation of their *R* scores (**Methods**). The relevance correlation might approximate phenotypic correlation since if two phenotypes are correlated, their relevance to human contexts will also be correlated. We used two GWAS datasets with phenotypic correlation computed from individual phenotypic data as the gold standard and compared SpecVar to two other methods LDSC-SAP and LDSC-SEG.

The first dataset was GWAS of 78 distances on the human face (Xiong et al., 2019). Based on summary statistics, we computed the relevance correlation of 3,003 pairs of distances with SpecVar, LDSC-SAP, and LDSC-SEG. We compared the relevance correlation with phenotypic correlation from individual phenotypic data and computed the Pearson coefficient correlation (PCC, **Methods**) to evaluate the performance of these three methods. SpecVar’s relevance correlation showed the best performance in approximating phenotypic correlation (**Fig. 5a, b**, PCC=0.522), which outperformed the other three methods: LDSC-SAP PCC=0.467 (**Fig. 5b**), LDSC-SEG PCC=0.405 (**Fig. 5b**). We also evaluated the ability to approximate the phenotypic correlation of highly correlated phenotypes. By setting the threshold of phenotypic correlation to be 0.4, we obtained the 363 highly correlated phenotype pairs of facial landmark distances and compared the three methods based on their performance on these pairs of phenotypes. We found SpecVar also performed best with PCC 0.467, which was the largest among the three methods: LDSC-SAP PCC=0.454, LDSC-SEG PCC=0.245 (**Fig. 5c**). We used the mean square error as a metric to evaluate the performance (**Methods**) and SpecVar was also the best among the three methods (**Fig. S2**).

**Fig. 5.**
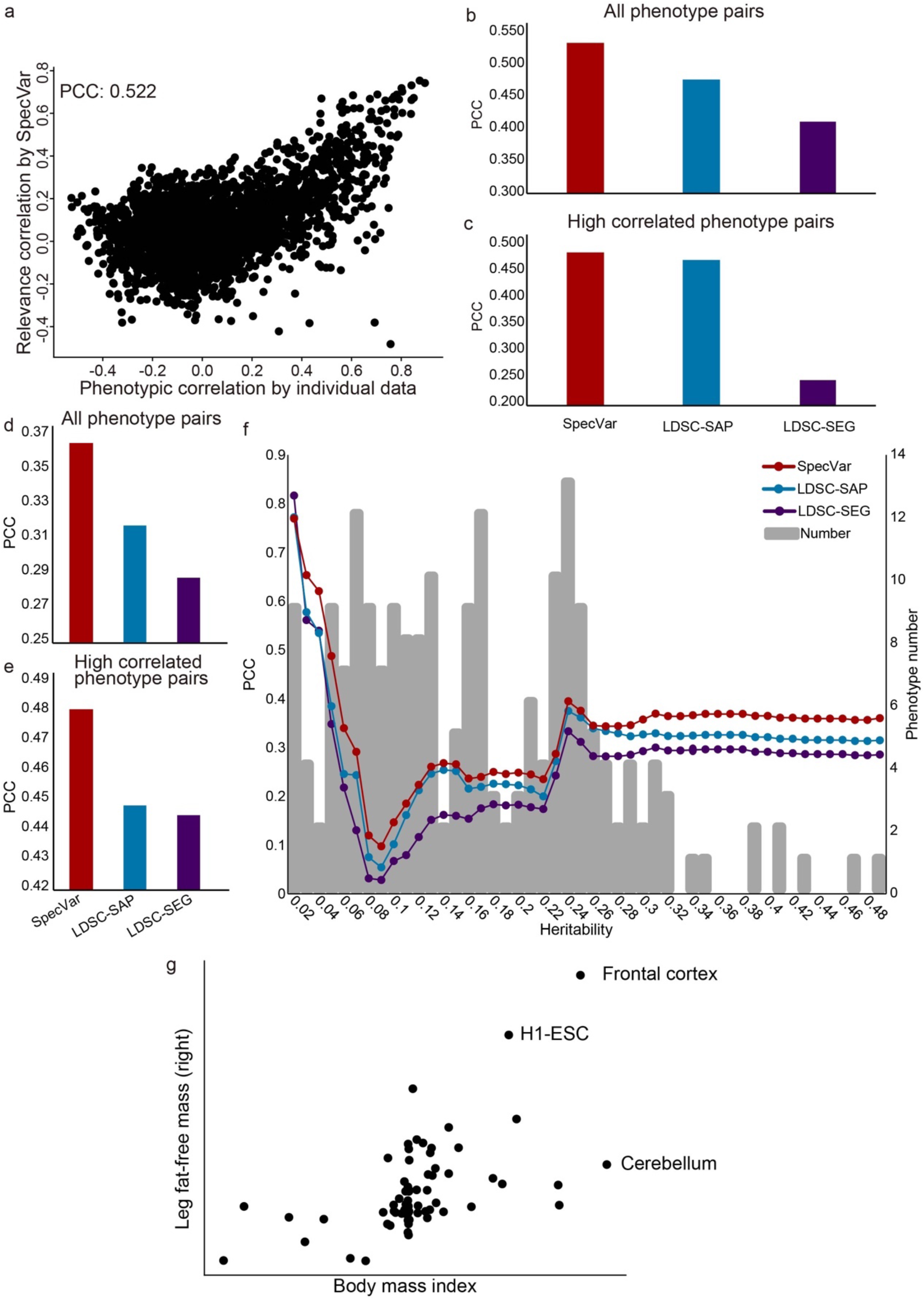
The scatter plot of true phenotypic correlation and relevance correlation by SpecVar. Each point means a pair of facial distances. (b) For all phenotype pairs of facial distances, the PCC between phenotypic correlation and relevance correlation of three methods. (c) For highly correlated phenotype pairs of facial distances, the PCC between phenotypic correlation and relevance correlation of three methods. (d) For all pairs of UKBB phenotypes, the PCC between phenotypic correlation and relevance correlation of three methods. (e) For highly correlated pairs of UKBB phenotypes, the PCC between phenotypic correlation and relevance correlation of three methods. (f) For UKBB phenotype pairs with different heritability thresholds, the PCC between phenotypic correlation and relevance correlation of four methods. (g) Scatter plot of *R* scores across 77 human contexts of body mass index and leg fat-free mass (right).

The second GWAS dataset was from UK-Biobank. There were 4,313 GWAS in UK-Biobank, from which we selected 206 high-quality GWAS summary statistics of 12 classes (**Table S3, Methods**). We applied SpecVar and the other two methods to obtain the relevance correlations among these 206 phenotypes and used the phenotypic correlation computed from individual data as validation. First, SpecVar performed best in the approximation of phenotypic correlation (PCC=0.360), followed by LDSC-SAP (PCC=0.315) and LDSC-SEG (PCC=0.285) (**Fig. 5d, Fig. S3a**). For highly correlated phenotypes, SpecVar’s relevance correlation was also closest to phenotypic correlation (**Fig. 5e, Fig. S3b**). We found that the heritability of these 206 phenotypes was quite variable. For example, “100630” (Rose wine intake) had a heritability of 6.52 × 10^−3^, and “5257_irnt” (Corneal resistance factor right) had a heritability of 0.336. So, we checked if the heritability would influence the quality of relevance correlation. To do this, we set different thresholds of heritability and obtained a subset of phenotypes for each threshold. Then for the phenotype subset of each heritability threshold, we computed the PCC between relevance correlation and phenotypic correlation. For almost all the thresholds of heritability, SpecVar showed the best performance of PCC (**Fig. 5f, Fig. S3c**), and the smallest variance regarded heritability among these three methods (**Fig. S3d, e**). This means that the relevance correlation of SpecVar could estimate phenotypic correlation more accurately and robustly. SpecVar can interpret the relevance correlation by the common relevant tissues and shared regulations of two phenotypes. For example, body mass index and leg fat-free mass (right) were correlated with a phenotypic correlation of 0.697. SpecVar obtained a relevance correlation of 0.602, while LDSC-SAP obtained a relevance correlation of 0.342 and LDSC-SEG obtained a relevance correlation of 0.437. SpecVar further revealed that these two phenotypes were correlated because they were both relevant to the “frontal cortex” (**Fig. 5g**). Body mass index has been reported to be related to frontal cortex development (Laurent et al., 2020) and relevant to the reduced and thin frontal cortex (Islam et al., 2018; Shaw et al., 2018). Obesity and fat accumulation are also revealed to be associated with the frontal cortex (Gluck et al., 2017; Kakoschke et al., 2019). SpecVar further extracted these two phenotypes’ SNP-associated regulatory networks in the “frontal cortex” and found their SNP-associated networks were significantly overlapped. The significant overlap was observed at SNP, RE, TG, and TF levels: *P* = 8.17 × 10^−63^ for SNPs, *P* = 1.38 × 10^−47^ for REs, *P* = 5.96 × 10^−25^ for TGs, and *P* = 8.23 × 10^−25^ for TFs (**Fig. S4**). The shared regulatory network was involved with body weight and obesity. For example, in the brain, *SH2B1* enhances leptin signaling and leptin’s anti-obesity action, which is associated with the regulation of energy balance, body weight, and glucose metabolism (Rui, 2014).

Through the application of relevance correlation to two datasets with the gold standard of phenotypic correlation, we concluded that SpecVar can use the accurate relevance score to define relevance correlation, which could better estimate phenotypic correlation and could reveal shared heritability with common relevant tissues and overlapped context-specific regulatory networks.

## Discussion

In this paper, we introduced the context-specific regulatory network, which integrated paired gene expression and chromatin accessibility data, to construct context-specific regulatory categories for better interpretation of GWAS data. SpecVar was developed as a tool to interpret genetic variants of GWAS summary statistics. The key message is that integrating chromatin accessibility and gene expression data into context-specific regulatory networks can provide better regulatory categories for heritability enrichment (Gazal et al., 2019). SpecVar is based on the popular model S-LDSC (Finucane et al., 2015), which includes 52 function categories as the baseline model. In addition, we showed extending the functional categories from non-context-specific regions to context-specific regions could improve the heritability enrichment, which is consistent with other studies based on gene expression (Hilary K. Finucane et al., 2018) and ChIP-seq (van de Geijn et al., 2020) data.

SpecVar outperformed the existing methods in three points. First, SpecVar defined relevance score based on both heritability enrichment and P-value. Because of the variability in the number of REs in the context-specific regulatory networks (**Table S4**), using only heritability enrichment or P-value will not give a stable estimation of the relevance of phenotype to tissues. For example, in the experiment of identifying six phenotypes’ relevant tissues, heritability could select most relevant tissues for LDL and TC to be the “right lobe of liver” but failed to get correct tissues for other phenotypes (**Fig. S5**). P-value could obtain correct tissues for CP (“frontal cortex”) and BrainShape (CNCC) but failed to get correct tissues for LDL, TC, EA, and Face (**Fig. S6**). By combining heritability enrichment and P-value into *R* score, SpecVar could prioritize correct relevant tissues for all the six phenotypes (**Fig. 3**). Like the *R* score-based relevance correlation, we could use the heritability enrichment and P-value to compute relevance correlation (**Fig. S7a, b**). We found heritability enrichment and P-value would give larger MSE (**Fig. S7c, e**) and lower PCC (**Fig. S7d, f**) than the *R* score, which showed that SpecVar’s *R* score can achieve a better approximation of phenotypic correlation. Those comparisons showed that the *R* score was a good metric to evaluate tissue’s relevance to the phenotype. Second, SpecVar’s regulatory categories had advantages over the existing functional categories to explain heritability. The context-specific regulatory networks formed regulatory categories enable better heritability enrichment than other methods (**Fig. 2**). The regulatory categories of SpecVar can be used to calculate *R* scores to identify relevant tissues more accurately than other methods (**Fig. 3**). And the *R* score of SpecVar can also be used to compute relevance correlation to better approximate phenotypic correlation than other methods when we do not have comprehensive phenotype measurement in each individual (**Fig. 5**). Third, with the constructed context-specific regulatory network atlas, SpecVar could further interpret the relevant tissue by SNP-associated regulatory networks (**Fig. 4**) and interpret relevance correlation by common relevant tissues and shared SNP-associated regulations in relevant tissues (**Fig. 5g, Fig. S4**). These three aspects made SpecVar an interpretable tool for heritability enrichment, identifying relevant tissues, and accessing associations of phenotypes.

Based on the accurate and highly interpretable relevant tissue identification, the relevance correlation of SpecVar provides us with another perspective of associations between two phenotypes: if two phenotypes are correlated, their relevance to human contexts will also be correlated. This rationale is independent of genetic correlation, which is the proportion of variance that two phenotypes share due to genetic causes and can be estimated with GWAS summary statistics by LDSC-GC (B. Bulik-Sullivan et al., 2015). When using measured phenotype value correlation as the gold standard of phenotype correlation, we found that SpecVar performed better when the heritability of phenotype was low while LDSC-GC performed better when the heritability was high (**Fig. S8a, b**). This indicated that the integration of relevance correlation and genetic correlation might give a better estimation of phenotypic correlation. We validated this idea by regressing phenotypic correlation on relevance correlation and genetic correlation in two GWAS datasets. For the phenotypes of facial distances, if we only use relevance correlation to regress phenotypic correlation, the coefficient of determination (R square) was 0.2720; if we only used genetic correlation, the R square was 0.0002; if we used the linear combination of relevance correlation and genetic correlation to regress phenotypic correlation, the R square was 0.2765, which was significantly higher than that only with SpecVar (F test of R square increase, *P* ≤ 1.77 × 10^−5^) or only with LDSC-GC (*P* ≤ 5.27 × 10^−213^); and if we used a product (non-linear combination) of relevance correlation and genetic correlation, the R square was much higher: 0.2911 (**Fig. S8c, d**). And for 206 phenotypes of UK-BioBank, if we only used relevance correlation, the R square was 0.1289; if we only used genetic correlation, the R square was 0.5614; if we used the linear combination of relevance correlation and genetic correlation to regress phenotypic correlation, the R square was 0.5927, which was significantly higher than that only with SpecVar (*P* ≤ 2.20 × 10^−16^) or only with LDSC-GC (*P* ≤ 2.20 × 10^−16^); and if we used a product of relevance correlation and genetic correlation, the R square was 0.7375, which was much improved (**Fig. S8e, f**). These results showed that relevance correlation and genetic correlation revealed the association of phenotypes in a complementary way.

Our work can be improved in several aspects. The usage of context-specific regulatory networks contributed most to the improvement of SpecVar. But the context-specific regulatory networks can only cover part of the regulatory elements and genetic variants, which are highly essential and representative. Higher-quality and more comprehensive regulatory networks will help obtain better representation. Currently, we built the atlas of regulatory networks of 77 human contexts and only included CNCC in the early developmental stage, which was far from complete. We expect more developmental stages will be included with multi-omics data from ENCODE (Consortium et al., 2020) and GTEx (Consortium, 2020). On the other hand, the 77 human contexts were tissues and cell lines. Single-cell-omics data (Han et al., 2020) will provide cell type level resolution and allows the extension of SpecVar to include broader cell types. The higher-quality and more comprehensive data will help SpecVar to construct better regulatory categories and improve interpretation. Lastly, it will be useful to extend the current approach using a model based on individual Whole Genome Sequencing data (Li et al., 2020).

## Methods

### Regulatory network inference with paired expression and chromatin accessibility data by PECA2

The regulatory networks were inferred by the PECA2 (Duren et al., 2020) model with paired expression and chromatin accessibility data. First, we collected paired expression and chromatin accessibility data of 76 human tissue or cell lines from ENCODE and ROADMAP (**Table S1**). Then with paired expression and accessibility data of each context, PECA2 calculated two scores. One was the trans-regulatory score. Specifically, PECA2 hypothesized that TF regulated the downstream TG by binding at REs. The trans-regulatory score was calculated by integrating multiple REs bound by a TF to regulate TG to quantify the regulatory strength of this TF on the TG. And PECA2 also considered a prior TF-TG correlation across external public data from ENCODE database. In detail, the TRS score *TRS*_*ij*_ of *i*-th TF and *j*-th TG was quantified as

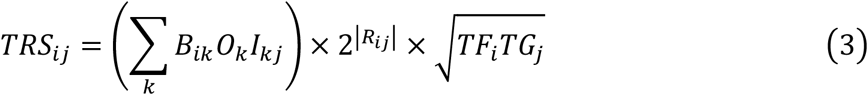

Here *TF*_*i*_ and *TG*_*j*_ were the expressions of the *i*-th TF and *j*-th TG. *B*_*ik*_ was the motif binding strength of *i-*th TF on *k-*th RE, which was defined as the sum of the binding strength of all the binding sites of *i -*th TF on *k-*th RE. *O*_*k*_ was the measure of accessibility for *k-*th RE. *I*_*kj*_ represented the interaction strength between *k*-th RE and *j*-th TG, which was learned from the PECA model on diverse ENCODE cellular contexts (Duren et al., 2017; Duren et al., 2018). *R*_*ij*_ was the expression correlation of *i*-th TF and *j*-th TG across diverse ENCODE samples. The significance of the TRS score was obtained by a background of randomly selected TF-TG pairs and the threshold of the TRS score was decided by controlling the false discovery rate (FDR) at 0.001.

The other one was the cis-regulatory score to measure the regulatory strength of RE on a TG. The cis-regulatory score *CRS*_*kj*_ of *k*-th RE on *j*-th TG was quantified as

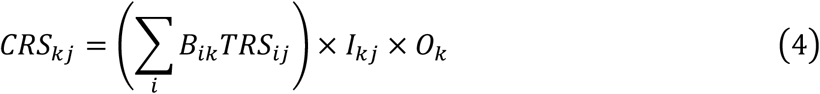

We approximated the distribution of log_2_+1 + *CRS*_*kj*_. by a normal distribution and predicted RE-TG associations by selecting the RE-TG pairs that have P-value≤0.05.

The output of PECA2 was a regulatory network with TFs, REs, and TGs as nodes and the regulations among them as edges. This procedure was applied to 76 human contexts with paired expression and chromatin accessibility data and obtained 76 regulatory networks. We noted that the regulatory network of early development stage CNCC was reconstructed recently (Feng et al., 2021) and we included the regulatory network of CNCC to form our regulatory network atlas of 77 human contexts.

### Construction of context-specific regulatory network atlas

The context-specific regulatory network was obtained based on the specificity of REs. In detail, we had 77 regulatory networks, and each regulatory network had a set of REs *RE*_*i*_, 1 ≤ *i* ≤ 77. Firstly, we hierarchically clustered 77 contexts’ the regulatory networks into 36 groups by trans-regulatory score (**Table S1**). Then for a given context, a RE was defined as a context-specific RE if it was not overlapped with REs of other contexts. Formally, the context-specific RE set of *i*-th context *C*_*i*_ was defined as

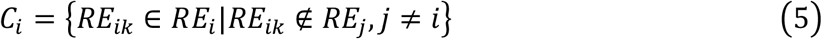

Here *RE*_*ik*_ ∉ *RE*_*j*_ means *RE*_*ik*_ was not overlapped with any REs in *RE*_*j*_:

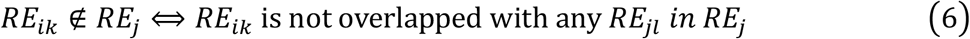

And we defined “overlapped” 1) for REs from contexts of the different groups, two REs were overlapped if their overlapping base ratio were over 50%; 2) for REs from contexts of the same group, two REs were overlapped if their overlapping base ratio were over 60%. The reason we used different “overlapped” criteria for REs from the same group and different groups was to retain group-specific REs. For example, for the brain tissues, we had five cell types: “Ammon’s horn”, “caudate nucleus”, “cerebellum”, “frontal cortex”, and “putamen”. If we defined RE’s specificity with stringent condition among these five brain cell types, many common brain REs would be lost.

Finally, the context-specific regulatory network was formed by specific REs and their directly linked upstream TFs and downstream TGs. And the context-specific RE sets *C*_*i*_, 1 ≤ *i* ≤ 77 gave the regulatory categories of SpecVar.

### Heritability enrichment and *R* score of GWAS summary statistics by SpecVar

SpecVar used stratified LDSC (Finucane et al., 2015) to compute partitioning heritability enrichment. Under the linear additive model, S-LDSC models the causal SNP effect on phenotype as drawn from a distribution with mean zero and variance

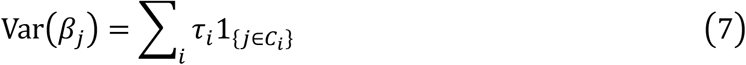

And with the assumption that the LD of a category that is enriched for heritability will increase the *χ*^2^ statistic of a SNP more than the LD of a category that does not contribute to heritability, the expected *χ*^2^ statistic is modeled as follows:

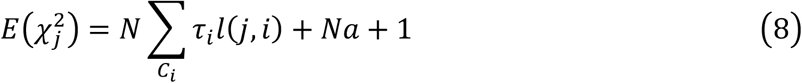

where *N* is the sample size, *C*_*i*_ denotes the regulatory category formed by the *i* -th context-specific regulatory network, 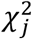 is the marginal association of SNP *j* from GWAS summary statistics, 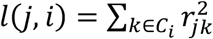 is the LD score of SNP *j* in the *i* -th category, *a* measures the contribution of confounding biases and *τ*_*i*_ represents heritability enrichment of SNPs in *C*_*i*_. S-LDSC estimates standard errors with a block jackknife and uses these standard errors to calculate the P-value *p*_*i*_ for the heritability enrichment (Finucane et al., 2015).

To make a trade-off between heritability enrichment score and P-value resulting from a hypothesis test, we combined heritability enrichment and statistical significance (P-value) to define the relevance score (*R*_*i*_) of this phenotype to *i*-th context as follows:

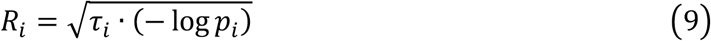

The relevance score (*R* score) offered a new robust means to rank and select relevant tissue for a given phenotype (Xiao et al., 2014) (**Fig. S5-7**).

### Four alternative methods to construct representations of GWAS summary statistics

Based on expression and chromatin accessibility data, there were four alternative methods for constructing regulatory categories: All Accessible Peaks (AAP), Specifically Accessible Peaks (SAP), Specifically Expressed Genes (SEG), and All Regulatory Elements (ARE).

The AAP method used all the chromatin accessible peaks of each context to form a genome functional category, which was used for partitioned heritability enrichment analysis. The SAP method used the same rules of SpecVar above to obtain context-specifically accessible peaks of each context, and the context-specific peaks sets of *M* contexts formed functional categories of SAP. The SEG method was constructed by following the procedure in (H. K. Finucane et al., 2018). First, the t-statistics for differential expression of each gene in each of the *M* contexts were calculated. Then for each context, the top 10% genes ranked by t-statistic were selected, and the 100Kb windows around those top 10% genes were used to form a functional category. For the ARE method, we obtained all REs in the regulatory network of a context to be a functional category, and the RE sets of *M* contexts formed regulatory categories of ARE.

We could conclude the relationship between the five methods: SpecVar, SAP, and SEG were methods based on specificity; SpecVar and SAP were based on the specificity of ARE and AAP, respectively. SpecVar and ARE used the expression and chromatin accessibility simultaneously; SAP and AAP only used the chromatin accessibility data; and SEG only used the gene expression data.

After obtaining functional categories with these four alternate methods, we could also use S-LDSC to obtain heritability enrichment and define the *R* score representation of GWAS summary statistics with equations (8) and (9). We called them LDSC-AAP, LDSC-SAP, LDSC-SEG, and LDSC-ARE, respectively. We compared these four alternate methods with SpecVar.

### Relevant tissue identification and relevance correlation analysis by SpecVar

SpecVar identified relevant tissues and defined relevance correlation based on *R* scores. The *R* scores to *M* contexts could be aggregated into a context-specific vector representation of GWAS summary statistics:

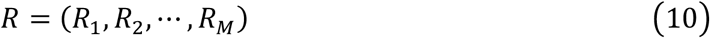

For a single phenotype, the *R* scores to *M* contexts could be used to get the relevant tissues. We used six phenotypes to analyze the distribution of *R* scores and found that the *R* scores followed a Gaussian distribution (**Fig. S9**).

We approximated the distribution of R to be Gaussian distribution and used the threshold P-value≤0.05 to get the relevant tissues. This gave the relevant tissues of the six phenotypes (**Table 1**), which was consistent with prior knowledge.

For two phenotypes, such as phenotype *p* and phenotype *q*, we obtained their *R* score representations:

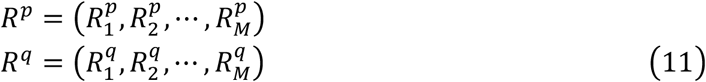

Then the Spearman correlation of their *R* score representation was used to define the relevance correlation:

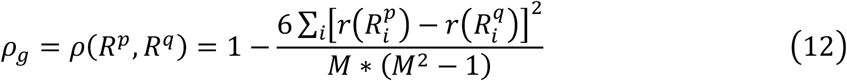

Here 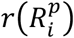 and 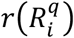were the ranks of *i* -th context by the relevance score for the two phenotypes.

For two other specificity-based regulatory categories LDSC-SAP and LDSC-SEG, we also used their functional categories to compute heritability enrichment and P-value and defined the *R* score with equation (7-9). The *R* scores of LDSC-SAP and LDSC-SEG were used to obtain relevant tissues and relevance correlation.

### Evaluation of relevant tissue identification and relevance correlation

To evaluate the performance of the SpecVar and other methods, we used different datasets as the gold standard.

For the application to identify relevant tissues, we used six well-studied phenotypes that we had knowledge of the relevant tissues: two lipid phenotypes (LDL and TC) were relevant to the liver; two human intelligential phenotypes (EA and CP) were relevant to the brain; two craniofacial bone phenotypes (Face and BrainShape) were relevant to CNCC. We used different methods to identify relevant tissues of these six phenotypes and checked if they obtained the correct tissues.

For relevance correlation, we used the phenotypic correlation computed with individual phenotypic data as the gold standard. First, we computed the Pearson correlation coefficient (PCC) between relevance correlation and phenotypic correlation:

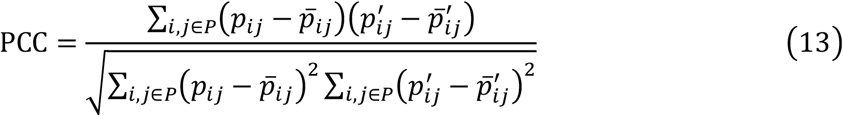

Here *P* was the set of phenotypes, and *N* was the number of phenotype pairs; *p*_*ij*_ was the phenotypic correlation computed with individual phenotypic data, and 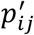 was the relevance correlation; 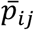 was the average of *p*_*ij*_, and 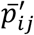 was the average of 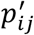. A larger PCC indicated better performance in approximating phenotypic correlation.

Another metric we used was the mean square error (MSE) between relevance correlation and phenotypic correlation:

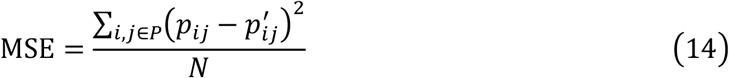

A smaller MSE indicated better performance in approximating phenotypic correlation.

### Extracting SNP-associated regulatory subnetworks in relevant tissues

Given a phenotype’s GWAS summary statistics and a context, SpecVar identified SNPs associated regulatory subnetwork by considering the following two factors: 1) the cis-regulatory score of SNP-associated RE should be large enough to indicate its importance in the regulatory network; 2) the risk signal of SNPs (i.e., P-value) on or near this RE should be large to indicate its association with phenotype. We combined these two factors to define the association score (*A* score) of SNP-associated REs.

First, the regulatory strength of *k*-th RE was measured by the maximum cis-regulatory score of this RE. Formally,

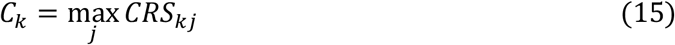

Here *CRS*_*kj*_ was the cis-regulatory score of *k*-th RE on *j*-th TG. For the *k*-th RE, the larger *C*_*k*_ was, the more important this RE was in the regulatory network. Second, the risk score of GWAS *S*_*k*_ for *k*-th RE was defined as the average of the -log(P-value) of SNPs located on or near this RE, which were down-weighted by their LD scores and distances to RE:

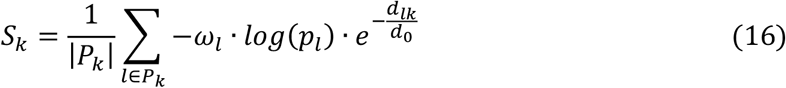

Here *P*_*k*_ was the set of SNPs whose distances were less than 50Kb to the *k*-th RE and |*P*_*k*_| was the total number of this SNP set; *ω*_*l*_ (the reciprocal of LD score, downloaded at https://data.broadinstitute.org/alkesgroup/LDSCORE/) was the weight of the *l*-th SNP; *p*_*l*_ was p- value of the *l*-th SNP in summary statistics; *d*_*lk*_ was the base pair distance of the *l*-th SNP to *k*-th RE and *d*_0_ was a constant, which was set to be 5,000 as default. For the *k*-th RE, a larger value of *S*_*k*_ indicated a stronger association with the given phenotype.

Finally, we obtained the association score (*A* score) of *k*-th RE by combining these two factors:

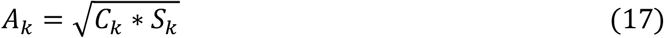

Every RE in the context-specific regulatory network was qualified by the *A* score. We used the GWAS of six phenotypes to analyze the distribution of *A* scores and found that the *A* scores followed a Gaussian distribution (**Fig. S10**). So, we hypothesized the distribution of *A* scores was Gaussian distribution and we selected the REs associated with the given phenotype by *A* scores’ FDR threshold of 0.05. The prioritized REs, as well as their directly linked upstream TFs, downstream TGs, and the associated SNPs, formed the SNP-associated regulatory subnetwork.

### GWAS summary statistics of UK-Biobank

The GWAS summary statistics of UK-Biobank were downloaded at http://www.nealelab.is/uk-biobank. There were 4,176 phenotypes and 11,372 GWAS summary statistics. We selected 206 GWAS summary statistics (**Table S3**) based on the following conditions.

1. Excluding sex-specific and “raw” type GWAS.
2. Sample size condition: *N* ≥ 50,000 and *N*_*control*_, *N*_*case*_ ≥ 10,000 for binary and categorical phenotypes.
3. Significant SNP number condition: the number of SNPs that pass the threshold of 5 × 10^−8^was not less than 500.
4. Manually curation: removing duplicated phenotypes, “job”, “parent” and “sibling” associated phenotypes.

## Data and code available

Codes and regulatory network resources are available at https://github.com/AMSSwanglab/SpecVar. Expression and chromatin accessibility data were summarized in Table S1. GWAS data used: GWAS summary statistics of LDL and TC were downloaded at http://csg.sph.umich.edu/willer/public/lipids2013/; GWAS summary statistics of EA (GCST006442), CP (GCST006572), BrainShape (GCST90012880-GCST90013164), and Face (GCST009464) were downloaded at GWAS catalog https://www.ebi.ac.uk/gwas/summary-statistics; GWAS summary statistics of UK-Biobank were downloaded at http://www.nealelab.is/uk-biobank. The LDSC genetic correlation and phenotypic correlation computed from individual phenotypic data were downloaded at https://ukbb-rg.hail.is/.

## Acknowledgments

We acknowledge funding from the National Key Research and Development Program of China (2020YFA0712402), Strategic Priority Research Program of the Chinese Academy of Sciences [XDPB17], and the National Natural Science Foundation of China (grants 12025107, 11871463, and 11688101).

## Author contributions

Y.W., and W.H.W. conceived and supervised the project. Z.F. designed the analytical approach and performed numerical experiments and data analysis. Z.D. contributed to the construction of regulatory networks. J.X., Q. Y., Y.H., B.S. contributed to interpreting biological insights. All authors wrote, revised, and contributed to the final manuscript.

---

**Fig. S1:**
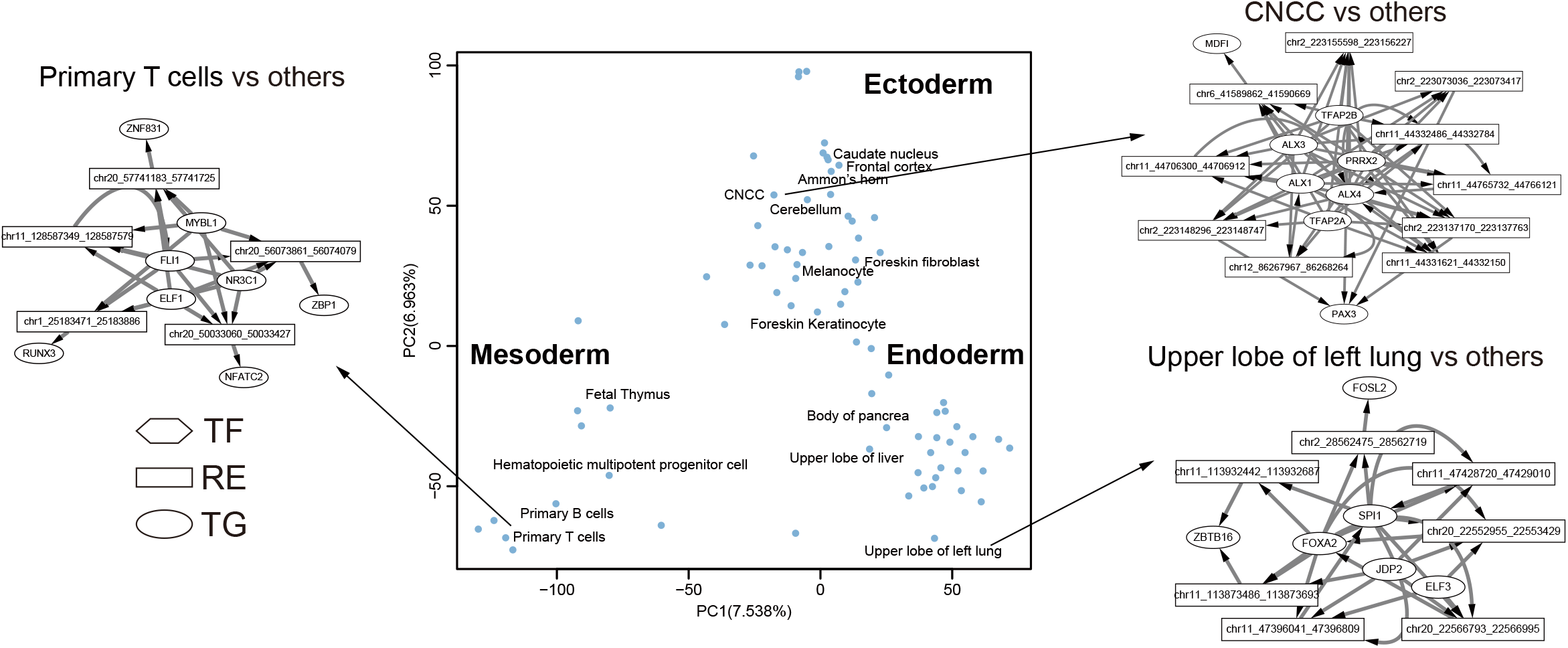
PCA plot of regulatory network atlas of 77 human tissues. The TRS score across 77 tissues are used for PCA analysis

**Fig. S2:**
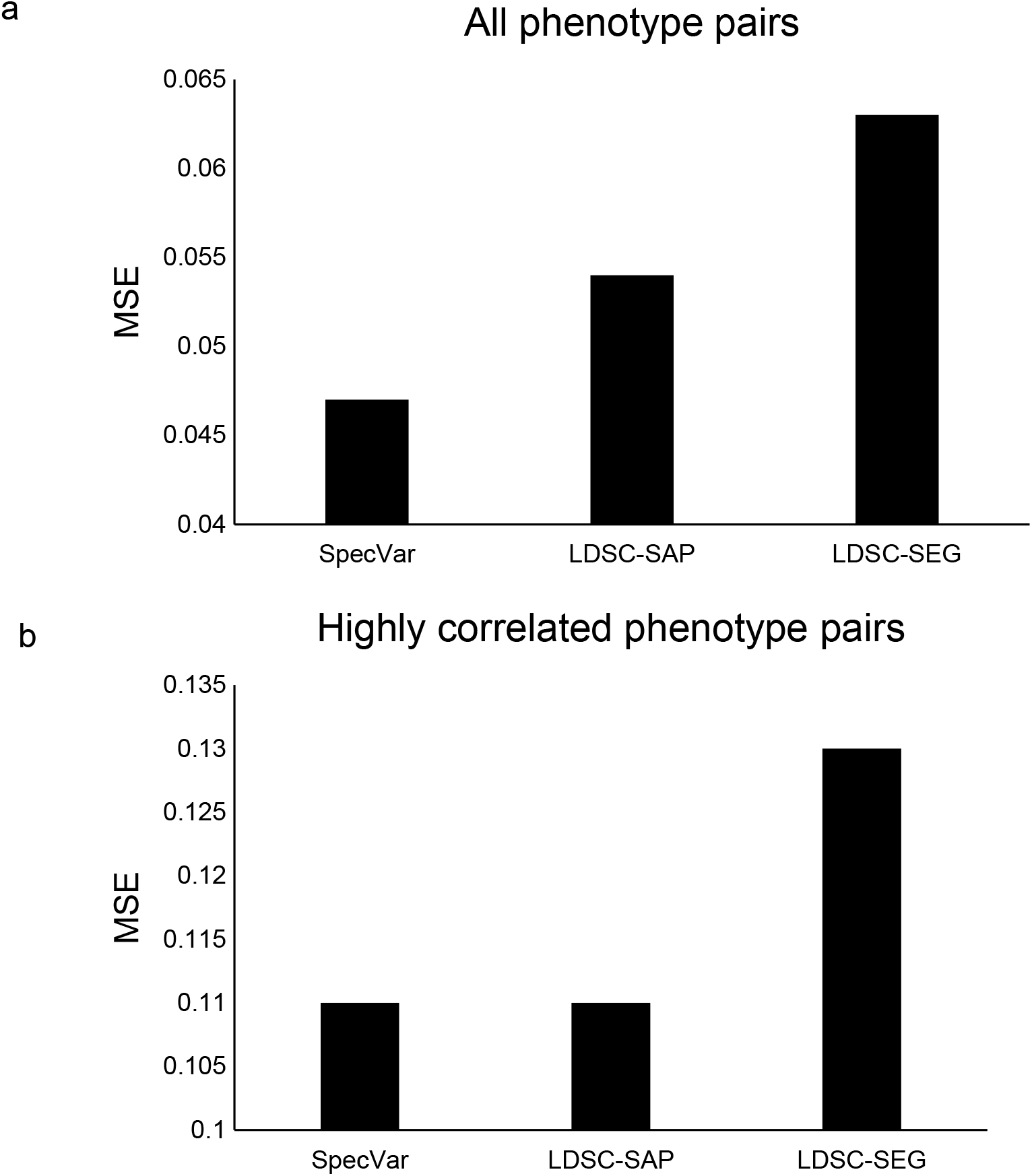
(s). For all phenotype pairs of facial distances, the MSE between true phenotypic correlation and relevance correlation of SpecVar, LDSC-SAP, and LDSC-SEG. (f). For highly correlated phenotype pairs of facial distances, the MSE between true phenotypic correlation and estimated phenotypic correlation of SpecVar, LDSC-SAP, and LDSC-SEG.

**Fig. S3:**
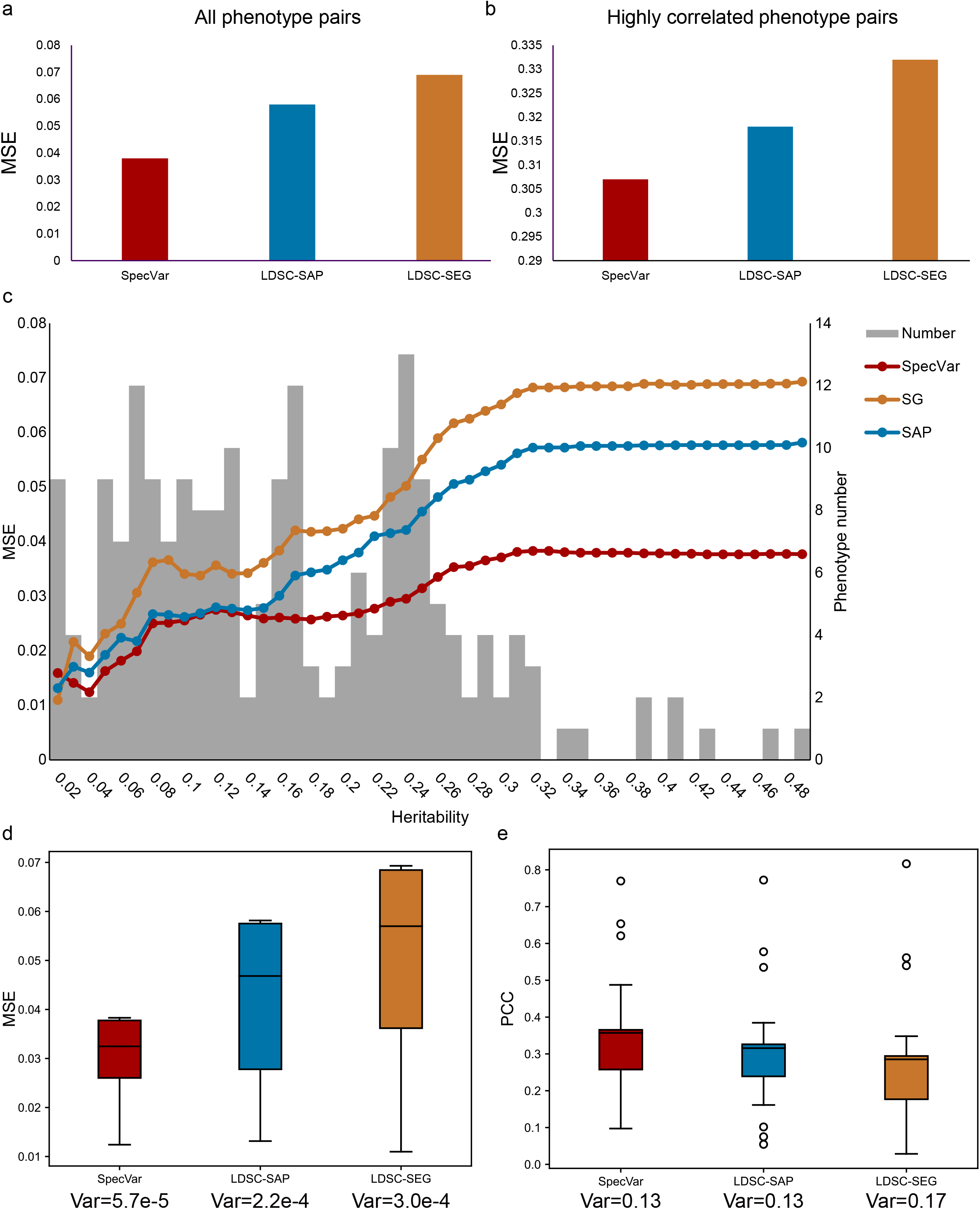
(a). For all phenotype pairs of facial distances, the MSE between true phenotypic correlation and relevance correlation of three methods. (b). For highly correlated phenotype pairs of facial distances, the MSE between true phenotypic correlation and relevance correlation of three methods. (c) For UKBB phenotype pairs with different heritability thresholds, the MSE between true phenotypic correlation and relevance correlation of three methods. (d). Boxplot of relevance correlation MSE under different threshold of phenotype heritability. Specvar shows the smallest variance. (e). Boxplot of relevance correlation PCC under different threshold of phenotype heritability. Specvar and LDSC-SAP shows the smallest variance.

**Fig. S4:**
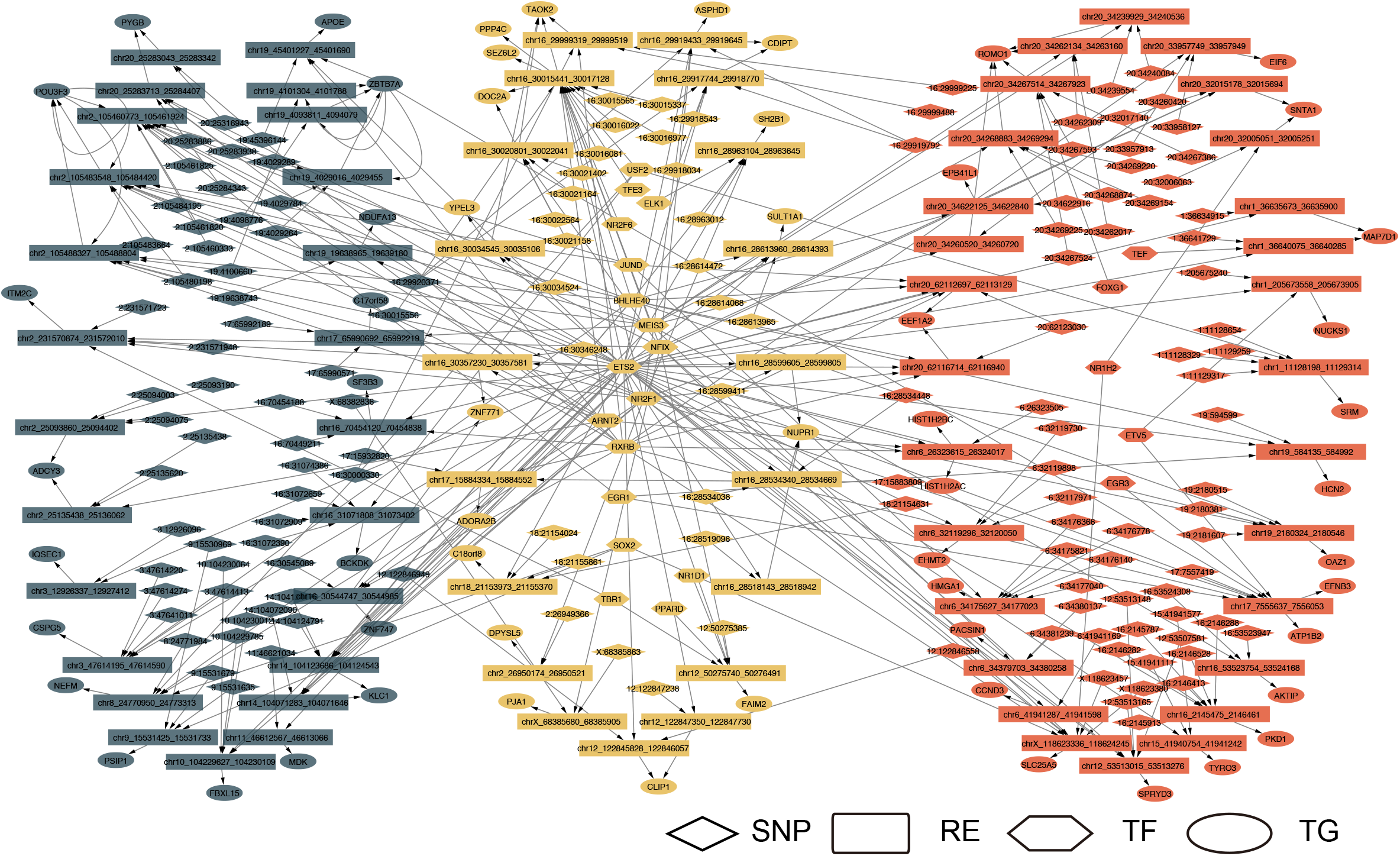
SNP associated regulatory network of “body mass index” (left) and “right leg fat-free mass” (right) in “frontal cortex”. These two phenotype associated regulatory networks are significantly overlapped (middle).

**Fig. S5:**
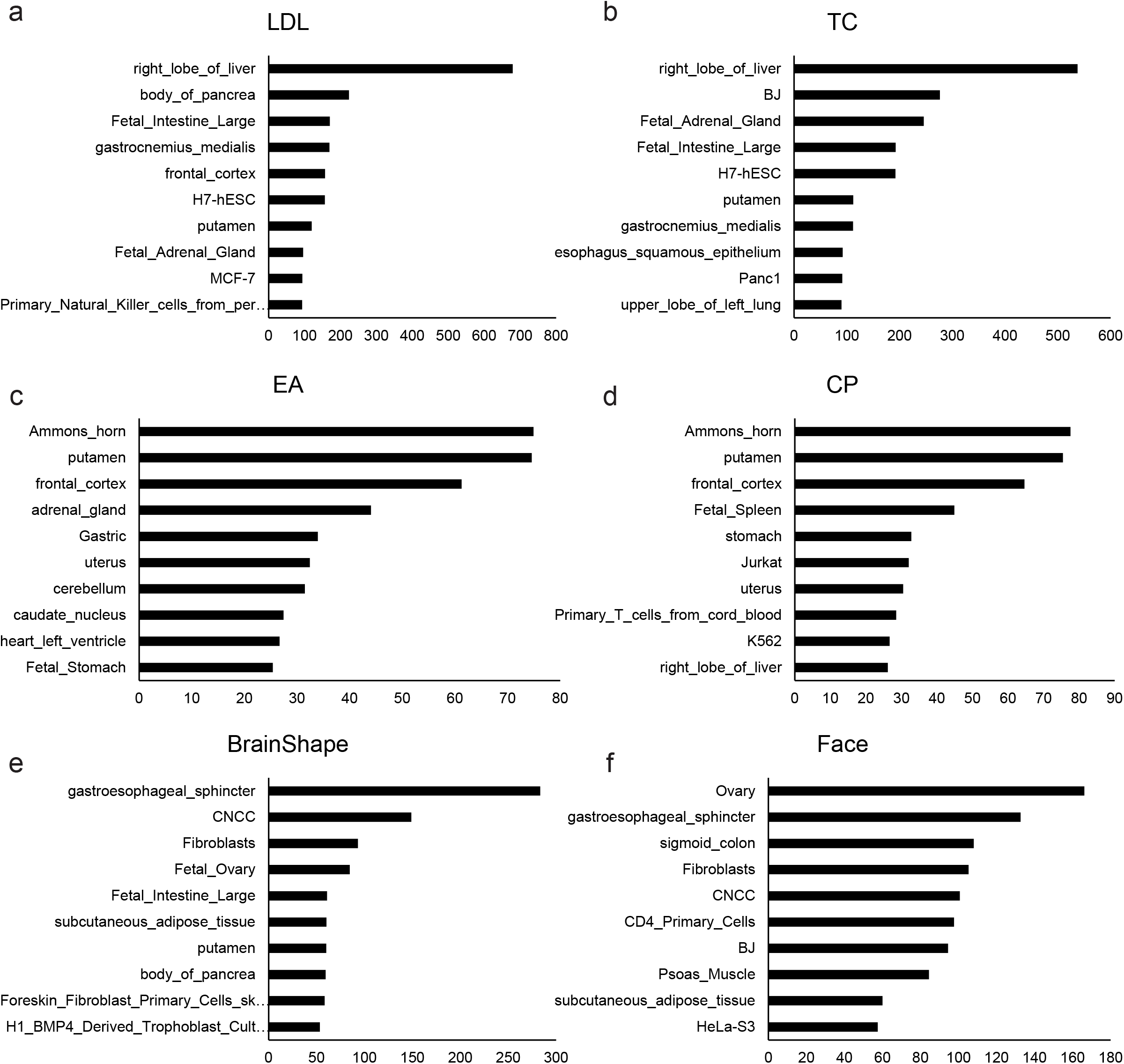
Top 10 contexts ranked by heritability enrichment in context-specific regulatory elements of (a). LDL, (b). TC, (c). EA, (d). CP, (e). BrainShape, (f). Face.

**Fig. S6:**
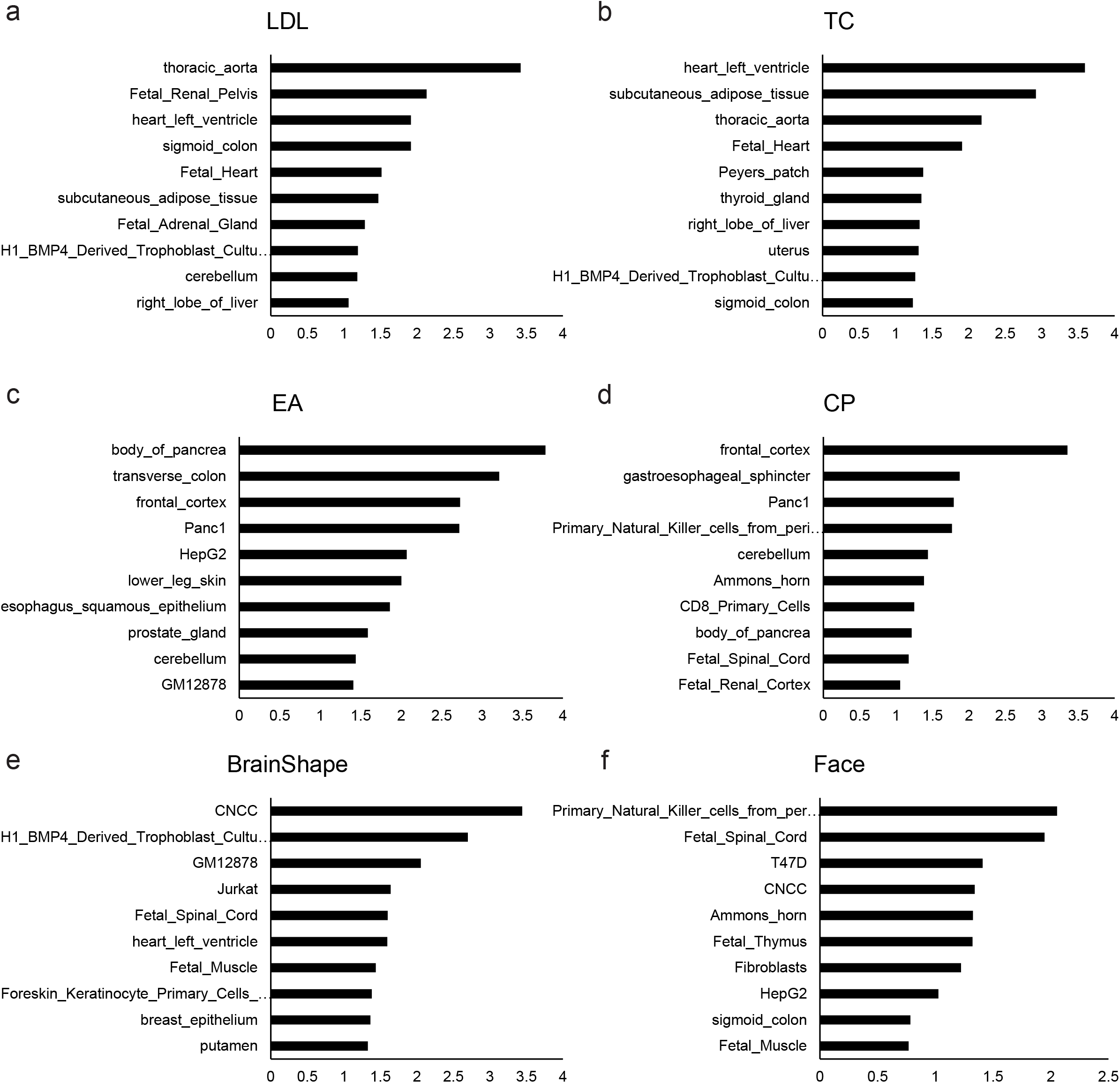
Top 10 contexts ranked by P-values of heritability enrichment in context-specific regulatory elements of (a). LDL, (b). TC, (c). EA, (d). CP, (e). BrainShape, (f). Face.

**Fig. S7.**
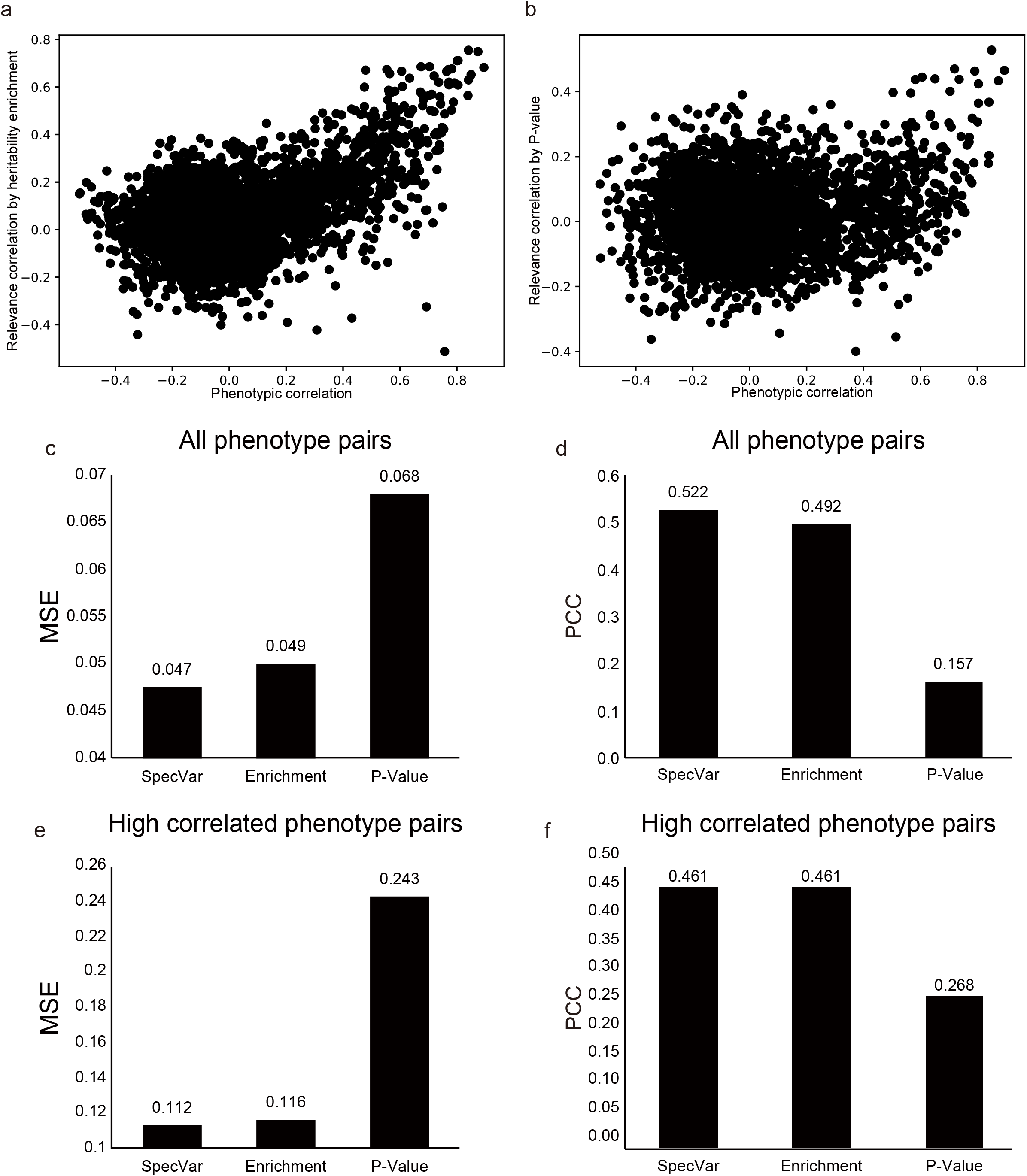
The scatter plot of true phenotypic correlation and estimated relevance correlation by (a). heritability enrichment, (b). -log(P-value). Each point means a pair of facial distances. (c). For all phenotype pairs, the MSE between phenotypic correlation and relevance correlation of SpecVar, heritability enrichment, and -log(P-value). (d). For all phenotype pairs, the PCC between phenotypic correlation and relevance correlation of SpecVar, heritability enrichment, and -log(P-value). (e). For high correlated phenotype pairs, the MSE between phenotypic correlation and relevance correlation of SpecVar, heritability enrichment, and -log(P-value). (f). For high correlated phenotype pairs, the PCC between phenotypic correlation and relevance correlation of SpecVar, heritability enrichment, and -log(P-value).

**Fig. S8.**
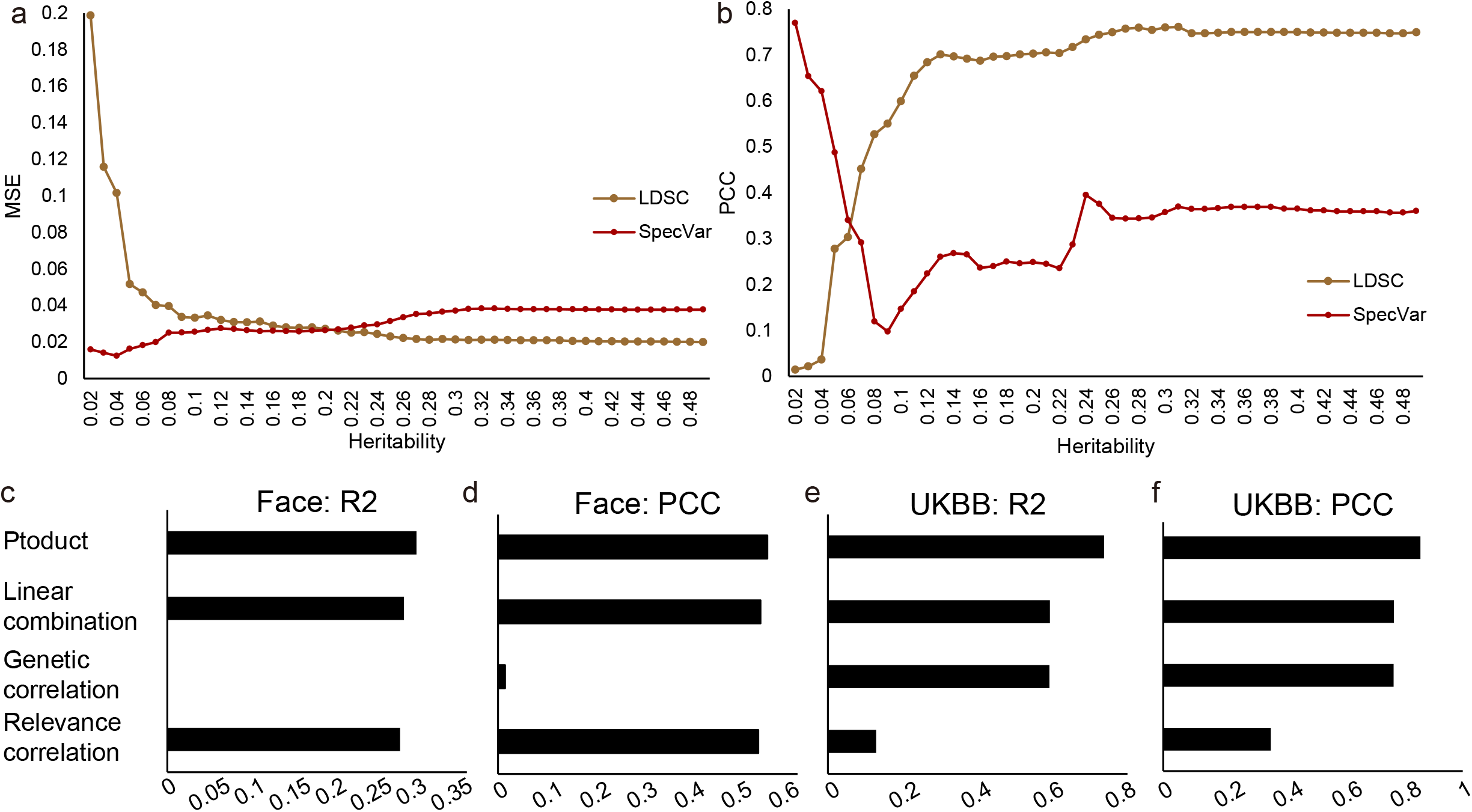
(a). For phenotype pairs with different heritability thresholds, the MSE between true phenotypic correlation and SpecVar’s relevance correlation; and MSE between true phenotypic correlation and LDSC-GC’s genetic correlation. (b). For phenotype pairs with different heritability thresholds, the PCC between true phenotypic correlation and SpecVar’s relevance correlation; and MSE between true phenotypic correlation and LDSC-GC’s genetic correlation. (c). The R2 metric of regression bertween phenotypic correlation and relevance correlation, genetic correlation, linear combination, product in Face distance phenotypes. (d). The PCC metric of four regression in Face distance phenotypes. (e). The R2 metric of four regression in UKBB phenotypes. (f). The PCC metric of four regression in UKBB phenotypes.

**Fig. S9.**
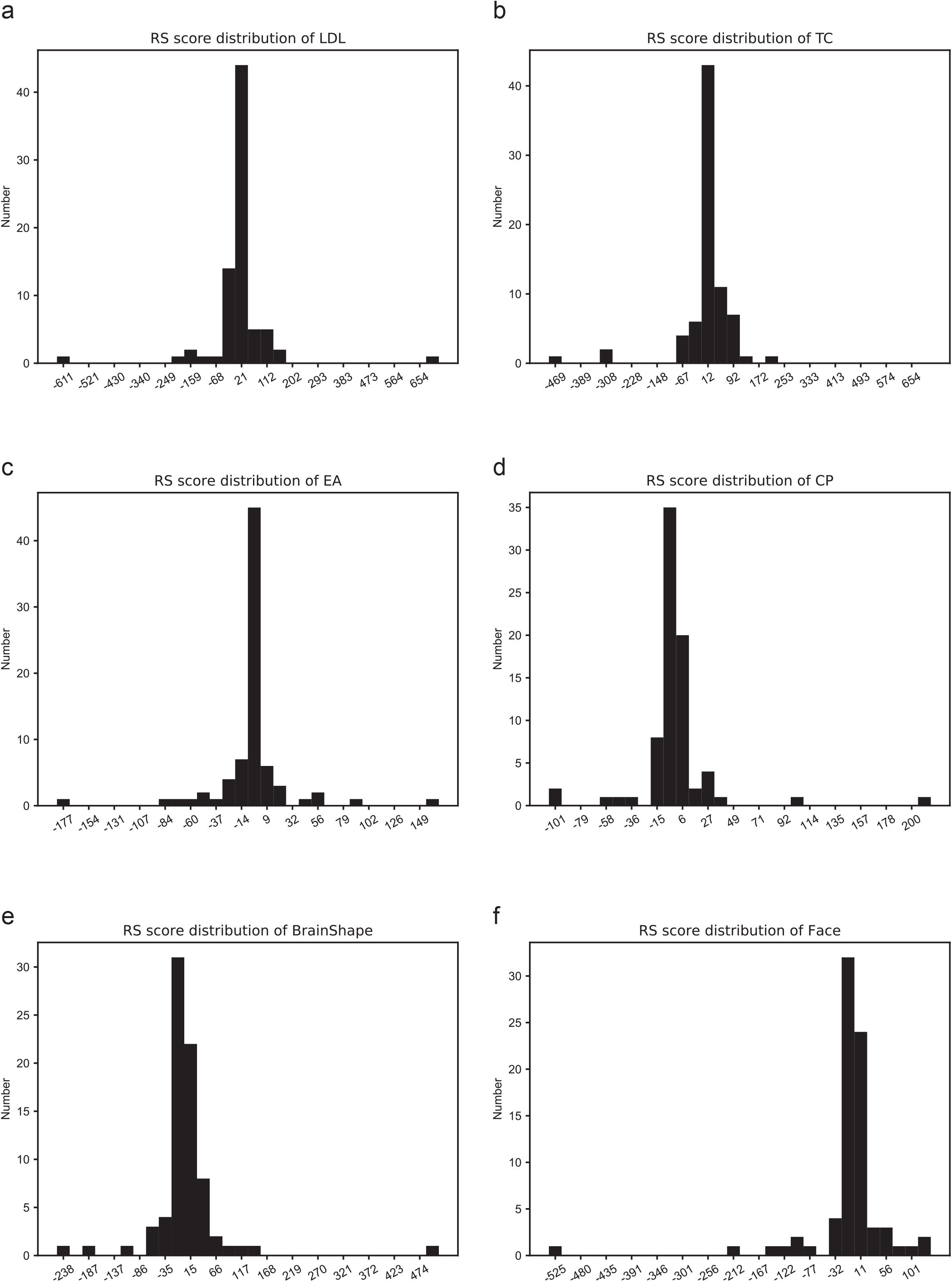
Distribution of *R* score of (a). LDL, (b). TC, (c). EA, (d). CP, (e). Face, (f). BrainShape.

**Fig. S10.**
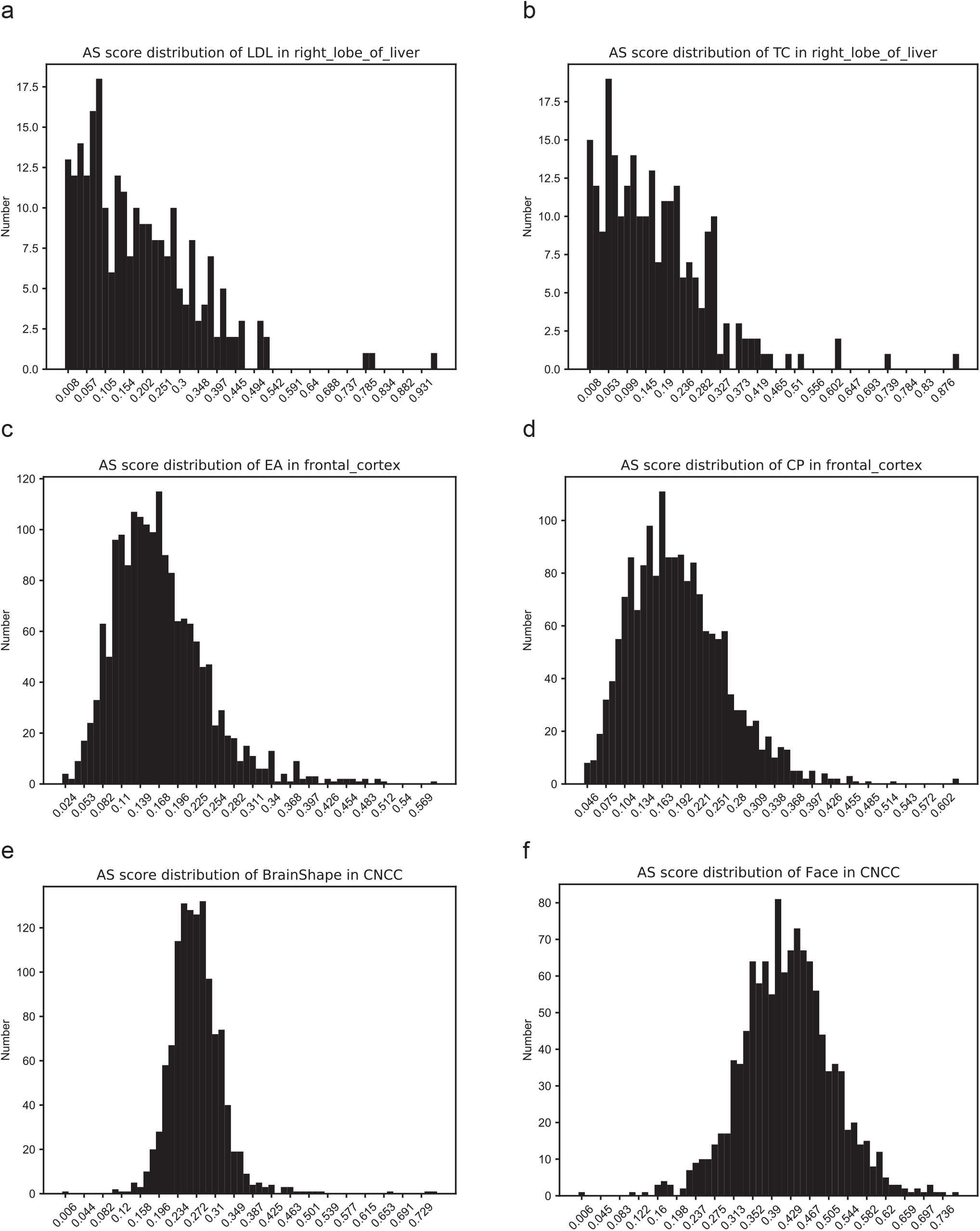
Distribution of *A* score of (a). LDL in “right lobe of liver”, (b). TC in “right lobe of liver”, (c). EA in “frontal cortex”, (d). CP in “frontal cortex”, (e). BrainShape in “CNCC”, (f). Face in “CNCC”.

